# Exploratory Network Analysis of Oral Bacteria Taste Signaling Autophagy Crosstalk in Oral Squamous Cell Carcinoma and Multi Target Ligand Design for the MAPK1 STAT3 mTOR Axis

**DOI:** 10.64898/2026.07.30.741861

**Authors:** Mahdi Akhavan, Hamid Laitifi-Navid, Amir Barzegar Behrooz, Sanaz Vakili, Rui Vitorino, Sajjad Aftabi, Sujatha Peela, SPD. Ponamgi, Robert J Schroth, Manuel Berumen, Cassandra Yuan, Tayebeh Akbari Azirani, Stevan Pecic, Prashen Chelikani, Saeid Ghavami

**Author notes:** Corresponding author: Saeid Ghavami, PhD, Department of Human Anatomy and Cell Science, College of Medicine, University of Manitoba, Lead contact: Saeid Ghavami, PhD, Department of Human Anatomy and Cell Science, College of Medicine, University of Manitoba, Winnipeg, Canada., Office Number. These authors have co-senior authorship (Stevan Pecic, Prashen Chelikani, Saeid Ghavami). These authors have co-first authorship (Mahdi Akhavan, Hamid Laitifi-Navid). These authors of co-second authorship (Amir Barzegar Behrooz, Sanaz Vakili).

## Abstract

G protein-coupled receptor (GPCR) signaling represents a critical interface between oral bacteria and host cellular regulation in oral squamous cell carcinoma (OSCC). Here, we integrated systems biology, exploratory machine learning, and structure-based drug design to characterize potential associations between bacteria-related signaling and autophagy and to identify candidate therapeutic targets. Taste-associated signaling genes belonging to the GPCR superfamily were curated from KEGG, while OSCC- and autophagy-associated proteins were obtained from STRING, Reactome, UniProt, KEGG, and HMDB. Ten bacteria-associated host-interaction datasets were integrated using NetworkAnalyst to construct protein– protein interaction networks, and key hub nodes were identified through degree and betweenness centrality. Feature matrices derived from network topology were analyzed using exploratory dimensionality reduction (PCA), hierarchical clustering, and supervised models (SVM and Gradient Boosting) to assess whether network-derived features showed separability according to literature-informed bacterial reference categories; a Dysbiosis Index was additionally calculated.

Results suggested that bacterial sensing through taste-associated GPCR signaling may converge on a MAPK1-centered axis linking calcium signaling, autophagy, and oncogenic pathways. Pathobiont-associated networks showed greater representation of inflammatory and terminal-autophagy-related signaling through MAPK1–STAT3, whereas commensal-associated networks were more closely aligned with cytoprotective autophagy through balanced MAPK1–TP53/PTEN networks. Exploratory machine learning analyses highlighted MDM2 and AKT3 as high-contribution, network-associated candidate features linked to group separability within the current dataset. A dual-target MTDL (SG101) was designed to target downstream nodes (MDM2 and JAK2), showing favorable predicted docking interactions and computationally predicted ADMET properties.

In conclusion, bacteria-associated host taste signaling may be linked to differing autophagy-related network states in OSCC, and targeting downstream regulatory hubs with multi-target ligands represents a hypothesis-generating strategy that warrants experimental validation for pathway-oriented therapy.

## 1. Introduction

Oral squamous cell carcinoma (OSCC) is the most common malignancy of the oral cavity and remains associated with poor survival, particularly in advanced and recurrent disease. Beyond classical carcinogens such as tobacco, alcohol, and human papillomavirus, dysbiosis of the oral microbiome has emerged as a key environmental driver that shapes epithelial transformation, chronic inflammation, and therapeutic resistance in OSCC [1]. Recent work indicates that oral bacteria modulate autophagy, apoptosis, and epithelial– mesenchymal transition (EMT) through oncogenic hubs, including MAPK1/ERK, STAT3, PI3K/AKT/mTOR, and TP53, positioning these pathways at the nexus of microbiome–tumor crosstalk [2–5].

Periodontal and OSCC-enriched taxa such as *Porphyromonas gingivalis*, *Fusobacterium nucleatum*, *Prevotella intermedia*, and *Peptostreptococcus stomatis* activate IL-6/STAT3, NF-κB, and PI3K/AKT/mTOR signaling, reprogramming autophagy to support tumor cell survival, invasion, and immune evasion [6]. Experimental and clinical studies link *P. gingivalis* and *F. nucleatum* to cyclin D1 upregulation, apoptosis suppression, and altered autophagic flux, consistent with network analyses in OSCC that converge on MAPK1, STAT3, CTNNB1, and EGFR as high-centrality nodes across multiple bacterial interaction networks [7]. Conversely, commensal and probiotic species such as *Lactiplantibacillus plantarum* and *Lactobacillus* spp. appear to preserve homeostatic autophagy, inhibit EMT, and buffer oxidative and metabolic stress, suggesting a protective role against malignant progression [2, 7, 8].

Oral microbes also communicate with host epithelia through taste receptor–like G protein–coupled receptor (GPCR) signaling, involving bitter, sweet and umami taste receptors (T2Rs or TAS2Rs, T1Rs or TAS1Rs) and their associated G protein subunits [9–12]. These receptors are broadly expressed in extra-oral tissues and regulate innate immunity, apoptosis, and barrier function, with bitter receptors exerting context-dependent pro-apoptotic or pro-survival effects [13–15]. In the oral cavity, bacteria-derived metabolites can engage GPCR/taste receptor pathways, altering calcium and cAMP signaling and feeding into ERK/STAT3 and mTOR signaling cascades, providing a route by which dysbiosis shapes dysgeusia and tumor behavior in OSCC [16–21].

Despite growing recognition of microbiome and autophagy involvement in OSCC, frameworks that link specific oral taxa, taste receptor–mediated GPCR signaling, and network-level autophagy reprogramming across the commensal pathobiont spectrum remain limited [4, 22]. This study synthesizes current evidence on the oral microbiome autophagy axis in OSCC, with a focus on taste receptor signaling, and introduces an exploratory and hypothesis-generating network-based model in which distinct microbes are predicted to converge on MAPK1/STAT3/mTOR hubs and may influence shifts in autophagy between tumor-promoting and cytoprotective states [3, 17, 23]. Understanding this microbiome–GPCR–autophagy triad may inform new prognostic and therapeutic strategies, including “autophagy-aware” targeting of MAPK1/STAT3/mTOR, modulation of taste receptor pathways, and probiotic approaches to restore homeostatic autophagy in OSCC.

Importantly, the convergence of microbial sensing, taste receptor–mediated GPCR signaling, calcium flux, and autophagy regulation highlights the inherently multi-pathway nature of OSCC biology. These interconnected signaling networks converge on central regulatory hubs such as MAPK1, STAT3, and PI3K/AKT/mTOR, which may collectively influence whether autophagy functions as a cytoprotective or tumor-promoting process. As a result, therapeutic strategies that target a single pathway are often insufficient to effectively control disease progression in such complex systems.

The traditional single-target approach with one drug is often limited for the treatment of complex diseases, which are driven via interconnected biological pathways [24–26]. Accordingly, single target-mediated drugs commonly fail to produce sufficient therapeutic efficacy. In a different approach, known as combination therapy, two or more drugs are used to simultaneously address different targets [27, 28]. While this approach enhances therapeutic effect, it is associated with limitations including drug–drug interactions, inconsistent pharmacokinetics, and reduced patient compliance [29–33]. A novel multi-target directed ligand (MTDL) strategy has therefore emerged, enabling the design of a single molecule capable of simultaneously targeting multiple biological components such as enzymes, receptors, and signaling pathways [34–39]. This approach allows for improved therapeutic efficacy, reduced adverse interactions, and a more coordinated pharmacological response.

In the context of cancer, including OSCC, where multiple signaling pathways govern proliferation, survival, metabolism, and stress adaptation, MTDL-based strategies offer a potentially valuable avenue. By simultaneously targeting interconnected nodes such as GPCR/taste receptors, calcium signaling pathways, autophagy regulators, and oncogenic hubs, MTDLs may provide an opportunity to address pathway redundancy and therapeutic resistance [40–43].

Understanding this bacterium–GPCR–autophagy triad therefore provides a hypothesis-generating framework for investigating OSCC progression and may offer a conceptual foundation for next-generation drug development. Targeting key signaling axes such as MAPK1/STAT3/mTOR, modulating taste receptor pathways, and restoring autophagy homeostasis may enable improved control of OSCC progression. In this context, multi-target drug design strategies, particularly MTDLs, represent a potential approach to simultaneously regulate bacterial signaling, intracellular stress pathways, and autophagy dynamics, offering a pathway-centric strategy that warrants further experimental and clinical investigation in OSCC.

## 2. Method and materials

### 2.1 Identification of Proteins Involved in the Taste Receptor Signaling Pathway

The identification of genes involved in the taste signaling pathway — Taste transduction (Homo sapiens [human]) — was conducted through a comprehensive review of the KEGG database (hsa04742)[44]. These genes encompass critical components, including G protein-coupled receptors, downstream effectors, the canonical PLC-β/IP□ signaling cascade, and ion channels essential for taste perception. This curated gene set provides a robust foundation for subsequent analyses and functional investigations into the correlations among taste transduction, the oral bacteria, autophagy, and oral squamous cell carcinoma. In total, 86 key genes implicated in this pathway were meticulously curated and are presented in **Supplementary Table 1**.

### 2.2 Proteomic Identification of Host Proteins Associated with Oral Squamous Cell Carcinoma

To curate a set of proteins implicated in oral squamous cell carcinoma (OSCC), we queried the STRING database (version 12.0; https://string-db.org/), a comprehensive resource for known and predicted protein– protein interactions derived from experimental data, curated databases, co-expression analyses, computational predictions, text mining, and orthology transfer[45]. Using OSCC-related disease terms and relevant pathway descriptors, we retrieved 125 proteins associated with OSCC biology or its signaling networks (**Supplementary Table 2**). This protein set served as the core OSCC-related host gene panel for subsequent comparative and integrative analyses.

### 2.3 Identification of Core Genes in the Autophagy Signaling Pathway

Genes involved in the autophagy signaling pathway were compiled by systematically querying multiple curated databases, including STRING (https://string-db.org/)[45], Reactome (https://reactome.org/)[46], UniProt (https://www.uniprot.org/)[47], KEGG (https://www.genome.jp/kegg/)[44], and HMDB (https://www.hmdb.ca/)[48]. Pathway identifiers and database accession numbers used are listed in **Table 1**. After integrating the retrieved data and removing duplicates (using official HGNC gene symbols as the unique identifier), a final list of 1,018 unique genes associated with the autophagy signaling pathway was obtained (**Supplementary Table 3**).

**Table 1.** Pathway identifiers and database accession numbers used for systematic retrieval of autophagy-related genes.

| **STRING** |  |  |  |
| --- | --- | --- | --- |
| GO:0006914 | Autophagy | GO:0044754 | Autolysosomes |
| GO:0016236 | Macroautophagy | GOCC:0061908 | Phagophore |
| GO:0044804 | Autophagy of the Nucleus | GO:0000407 | Phagophore Assembly Site |
| GO:0005776 | Autophagosome | GO:0034497 | Protein Localization to Phagophore Assembly Site |
| GO:0061912 | Selective Autophagy | GO:0034045 | Phagophore assembly site membrane |
| GO:0000045 | Autophagosome Assembly | GO:0097632 | Extrinsic component of phagophore |
| GOCC:0005776 | Autophagosome | GOCC:0097632 | Phagophore |
| **Reactome** |  | **KEGG** |  |
| R-HSA-9612973 | Autophagy | hsa04140 | *Autophagy* (or macroautophagy) |
| R-HSA-1632852 | Macroautophagy | hsa04136 | Autophagy - other - Homo sapiens (human) |
| **UNIPROT** |  |  |  |
| KW-0072 | Autophagy |  |  |

### 2.4 Identification of Host Protein Targets Modulated by Oral Bacteria in Oral Squamous Cell Carcinoma

To identify host protein targets acting as key effectors in oral bacteria-driven oncogenic signaling in oral squamous cell carcinoma (OSCC), along with their associated downstream pathways, we conducted a systematic review of the relevant literature [22, 49]. Ten clinically and experimentally prioritized oral bacteria taxa, together with their reported host protein targets and proposed mechanistic links to OSCC, were curated and categorized. Subsequently, separate protein–protein interaction (PPI) networks were constructed using NetworkAnalyst [50] for (a) crucial key proteins associated with the Taste Receptor Signaling Pathway, (b) high-priority OSCC-associated genes, and (c) genes comprising the canonical autophagy signaling pathway. Hub nodes within each network were identified based on Degree and Betweenness Centrality metrics [51, 52]. By integrating (i) hub Proteins associated with Taste Receptor Signaling Pathways, (ii) OSCC-derived hub proteins, (iii) autophagy pathway hub proteins, and (iv) literature-curated host targets for each of the ten bacterialtaxa, we generated ten bactiaa-centric candidate protein sets. Finally, taxon-specific PPI networks were built for each bacterial species using NetworkAnalyst, with network visualization and layout performed via the Steiner Forest algorithm to delineate the core backbone of each interactome (**Supplementary Tables 4–23**).

### 2.5 Data Acquisition and Feature Matrix Construction

The computational analysis was based on ten PPI networks, each representing the literature-supported, putative influence of a specific oral bacterial species on the human host proteome. For each network, raw data were provided in Microsoft Excel (.xlsx) format, containing gene identifiers (“Label”) and their corresponding Degree centrality values.

All analyses were performed in Python 3, using the pandas library for data handling. First, all ten network files were systematically loaded. To generate a unified feature space, Degree centrality values for all unique genes across the ten networks were aggregated into a single matrix. Degree centrality was selected as the primary feature due to its direct interpretability as a measure of node connectivity and relative topological prominence within the reconstructed bacteria-perturbed networks.

A comprehensive feature matrix of size 10 × N g was constructed, where each row represented one microbial species (analytical network unit) and each column represented a unique host gene (feature). Missing values—indicating that a gene was not represented in a specific reconstructed network under the applied network-construction criteria—were imputed with zero. Accordingly, a zero value was interpreted as topological non-representation in the corresponding network rather than evidence of biological absence, lack of expression, or lack of functional relevance.

Each bacterial species was assigned to one of three literature-informed reference categories based on established biological evidence: Pro-cancer, Anti-cancer, or Intermediate. These reference categories were used for exploratory supervised analyses and were not considered independently observed ground-truth classes.

### 2.6 Unsupervised Exploratory Analysis

To explore the intrinsic structure of the high-dimensional feature space, two unsupervised methods were applied using scikit-learn and scipy libraries.

Principal Component Analysis (PCA) was performed following Z-score normalization of the feature matrix. An initial PCA included all ten bacterial species. Subsequently, a secondary PCA was conducted after exclusion of two dominant topological profiles (Fusobacterium nucleatum and Streptococcus mitis) to visualize lower-magnitude variation and improve the resolution of patterns among the remaining species. These two species were not considered erroneous observations or removed from the overall biological interpretation; the secondary analysis was used only as a complementary visualization of structure that was less apparent in the complete ten-network PCA.

In parallel, hierarchical clustering was applied using Ward’s linkage method, which minimizes within-cluster variance. Euclidean distance was used as the similarity metric, providing a complementary visualization of similarity among the PCA-derived network profiles rather than an independent validation of the groupings.

### 2.7 Statistical Assessment of Clustering Stability

The stability and non-random structure of observed clustering patterns within the current ten-network dataset were assessed using a permutation test (n = 10,000). The variance explained by the first two principal components in the real dataset (V_real) was compared against a null distribution generated by random shuffling of the feature matrix.

The empirical p-value was calculated as:

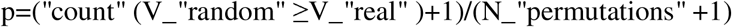

This analysis assessed whether the observed separation between bacterial reference categories could plausibly arise by chance under the specified null distribution. Given the limited number of bacterium-specific networks, the permutation results were interpreted cautiously as supportive evidence of structure within this dataset rather than proof of generalizability or biological validation.

### 2.8 Exploratory Supervised Analysis

Supervised models were used as exploratory tools to determine whether network-topological features contained sufficient structure to recover the literature-informed reference categories. These analyses were not designed to develop a diagnostic classifier or estimate performance in unseen clinical samples. Rather, they were used to identify candidate genes contributing to separation of the reconstructed taxon-specific networks.

A linear Support Vector Machine (SVM) (C = 1.0) was used to evaluate linear separability between classes. In addition, a Gradient Boosting Machine (GBM) classifier (n_estimators = 100, learning_rate = 0.1, max_depth = 3) was trained using a simple stratified split (70/30) for illustrative purposes. Because the dataset contained only ten bacterium-specific networks, the resulting performance estimates were considered highly sensitive to sample allocation and were not interpreted as reliable measures of out-of-sample predictive performance. Feature importance scores from the GBM model were used to identify genes showing relatively high contributions to class separation within this specific model. These estimates were interpreted as indicative, hypothesis-generating feature rankings rather than definitive biological drivers.

### 2.9 Pathway and *Candidate* Signature Analysis

Two candidate, hypothesis-generating functional gene signatures were defined based on PCA loadings. For each signature, a pathway activity score was calculated by summing Degree values of constituent genes. Scores were normalized to a 0–1 range using MinMaxScaler for visualization. Because these signatures were derived from the same ten-network dataset used for the exploratory analyses, they were not considered independently validated biological signatures.

A composite Dysbiosis Index was calculated as:

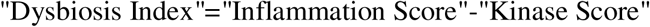

Statistical differences between bacterial reference categories were evaluated using a one-sided Mann– Whitney U test. The resulting differences were interpreted as exploratory associations within the current network dataset rather than predictive or clinically validated group distinctions.

#### Design of OSCC MTDL and ADMET Predictions

To design a MTDL capable of simultaneously promoting anti-cancer pathways and suppressing pro-cancer signaling in OSCC, we explored a pharmacophore-merging strategy based on two therapeutically relevant agents: idasanutlin and ruxolitinib. Idasanutlin, an investigational drug currently in Phase III clinical trials toward FDA approval, is a potent and selective MDM2 antagonist that reactivates the p53 tumor-suppressor pathway by disrupting the MDM2–p53 interaction, thereby inducing cell-cycle arrest and apoptosis (Table 2). Importantly, its oral bioavailability further supports its relevance for OSCC treatment.

**Table 2.** ADMET properties of Idasanutlin, Ruxolitinib and SG101.

|  | MW | LogP | nof<br>HBA | nof<br>HBD | BBB<br>Score | t<sub>1/2</sub> human<br>microsome (h) | t<sub>1/2</sub> human<br>plasma (h) | hERG<br>Inhibition | LogLD<sub>50</sub> |
| --- | --- | --- | --- | --- | --- | --- | --- | --- | --- |
| Idasanutlin | 616.48 | 8.0 | 8 | 3 | 1.39 | 0.38 | 3.34 | 0.06 | 1.66 |
| Ruxolitinib | 306.37 | 1.89 | 4 | 1 | 3.18 | 0.66 | 1.81 | 0.09 | 1.76 |
| SG101 | 545.91 | 4.47 | 10 | 3 | 0.35 | 0.70 | 4.35 | 0.01 | 1.62 |

Ruxolitinib, an **FDA-approved** JAK1/2 inhibitor, targets upstream regulators of STAT3 signaling, a central pathway driving tumor cell survival, proliferation, and immune evasion. In OSCC-relevant contexts, JAK– STAT3 signaling is frequently activated downstream of inflammatory cues such as IL-6. As highlighted in our network analysis later in text (Table 3), pro-cancer bacteria taxa including *Fusobacterium nucleatum* and *Porphyromonas gingivalis* converge on MAPK1–STAT3-centered inflammatory and autophagy-disruptive networks. Notably, *F. nucleatum* drives terminal inflammatory autophagy through a MAPK1–AKT–STAT3 axis, whereas *P. gingivalis* induces autophagy collapse via MAPK1–STAT3–NF-κB signaling, collectively reinforcing a strong rationale for targeting these pathways in OSCC.

**Table 3.**
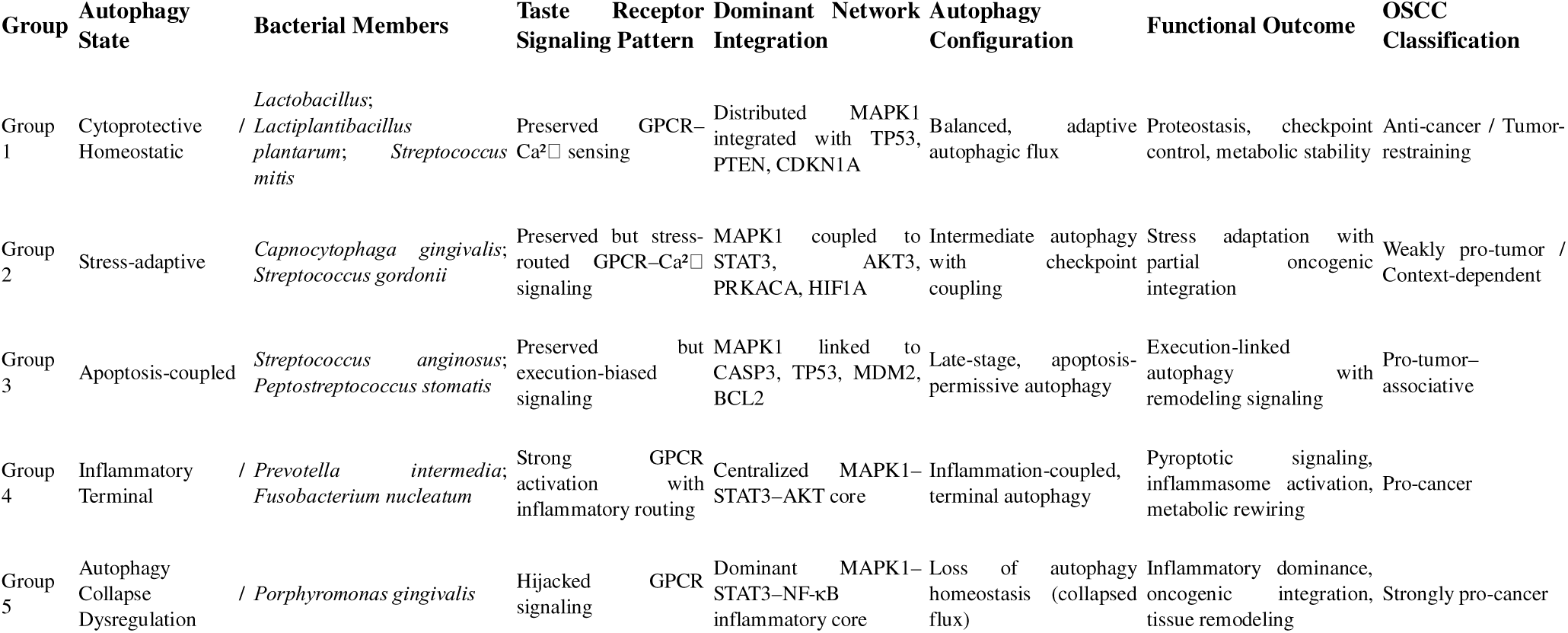
Stratification of bacterial taxa based on taste receptor–autophagy signaling continuum in OSCC.

Utilizing a merged pharmacophore approach [53–56], we designed SG101 by integrating key structural elements from both compounds. The nitrile group, present in both parent molecules, was retained due to its favorable electronic properties and contribution to binding interactions and pharmacokinetic optimization. The pyrazole core from ruxolitinib was selected over the pyrrolidine moiety of idasanutlin due to its critical role in maintaining potency and its advantageous planar structure, which simplifies synthesis by avoiding stereochemical complexity. The pyrrolo-pyrimidine scaffold, which mimics the adenine moiety of ATP and mediates JAK binding, was preserved to maintain kinase inhibitory activity. From idasanutlin, the 2-fluoro-3-chlorophenyl group was incorporated to support MDM2 binding and enhance metabolic stability, while the para-benzoic acid side chain was retained to improve solubility, polarity, and overall pharmacokinetic properties (Scheme 1).

**Scheme 1.**
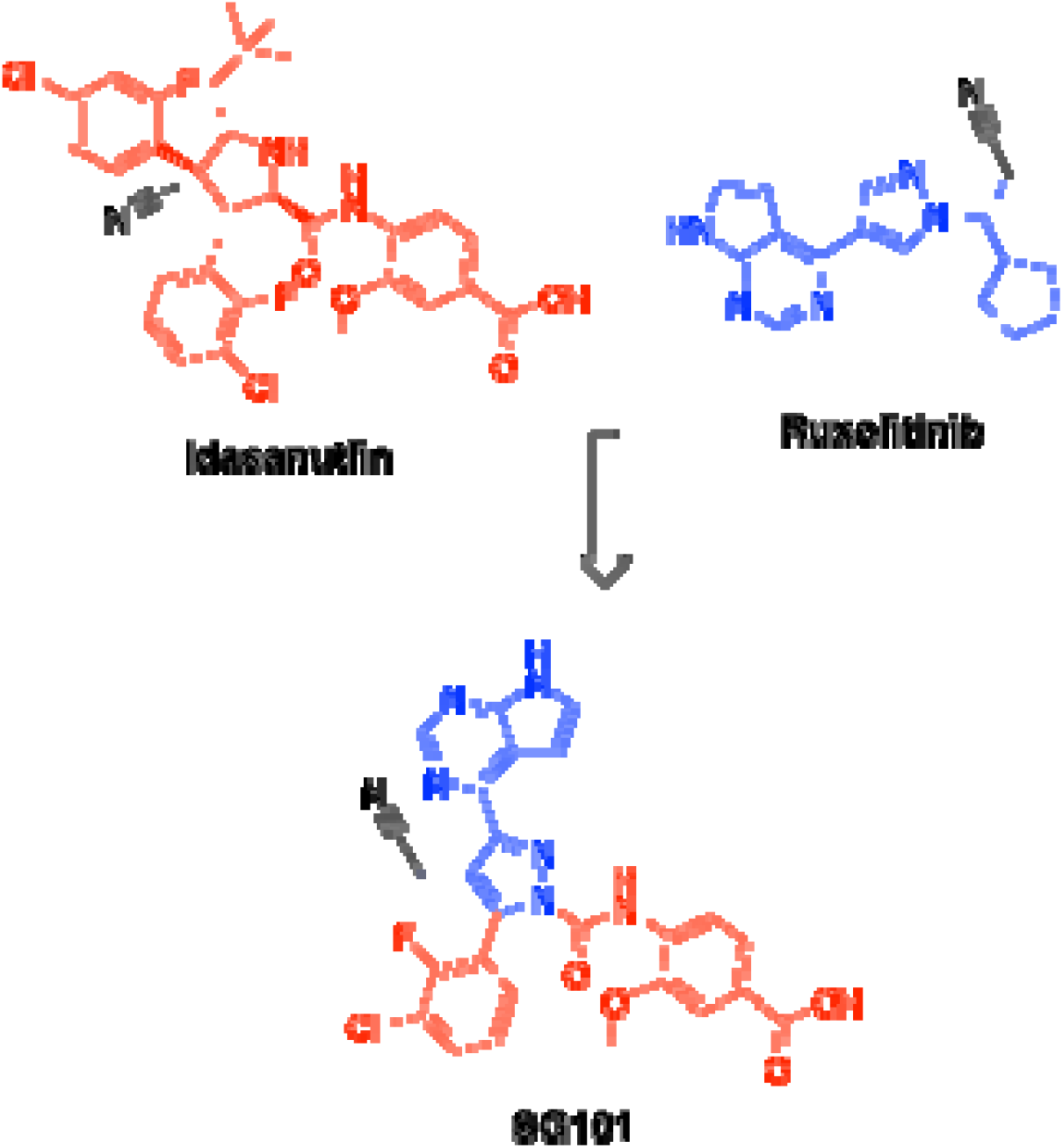
Design strategy for MTDL SG101: Idasanutlin (shown in red) is merged with Ruxolitinib (shown in blue). The common nitrile group is shown in black.

[ufig1]

We next performed in silico evaluation of Absorption, Distribution, Metabolism, Excretion, and Toxicity (ADMET) properties for SG101 and compared these with idasanutlin and ruxolitinib. Drug-likeness was first assessed using Lipinski’s Rule of Five [57]. As summarized in **Table 2**, ruxolitinib fully complies with all criteria (MW 306.37, LogP 1.89, HBA 4, HBD 1), consistent with favorable oral bioavailability. In contrast, idasanutlin violates two parameters (MW 616.48 and LogP 8.0), suggesting potential limitations in solubility and pharmacokinetic variability. SG101 shows a single violation (MW 545.91) while maintaining acceptable lipophilicity (LogP 4.47) and hydrogen bonding (HBA 10, HBD 3), indicating that it remains within a developable oral drug space despite its larger size.

Blood–brain barrier (BBB) permeability was evaluated to assess off-target CNS exposure. Based on established thresholds (BBB score > 4) [58], all compounds, including SG101 (BBB score 0.37), are predicted to have minimal brain penetration, which is desirable for OSCC-targeted therapy. These results are also summarized in Table 2.

Metabolic stability was assessed through predicted half-lives in liver microsomes and human plasma. Considering the impact of first-pass metabolism on orally administered drugs [59], all compounds exhibited relatively short microsomal half-lives, with SG101 showing comparatively improved stability. Notably, SG101 demonstrated enhanced plasma stability, suggesting reduced susceptibility to rapid hydrolysis and improved systemic persistence (Table 2).

Toxicity profiling included prediction of hERG inhibition and acute toxicity (logLD_50_). As shown in Table 2, all compounds exhibited low predicted risk of hERG channel inhibition (<0.5) [60] and logLD_50_ values between 1.5 and 2, indicating low-to-moderate acute toxicity. Collectively, these data support SG101 as a pharmacokinetically favorable and safe candidate for further preclinical evaluation.

## 3. Results

### Commensal-associated networks predict coupling of taste signalling to cytoprotective autophagy in OSCC

KEGG-based host–bacteria interactome analysis revealed that commensal-associated taxa, including *Lactobacillus*, *Lactiplantibacillus plantarum*, and *Streptococcus mitis*, establish a conserved signaling architecture in which epithelial taste receptor– GPCR signaling is preserved and coupled to adaptive, cytoprotective autophagy rather than inflammatory or lytic responses (**Figure 1**). Across all three taxa, upstream chemosensory modules composed of GNAS/GNAI1, GNB1/3, GNG2/13, and taste-linked receptors (GABBR1/2, GRM1) formed coherent detection systems, indicating active engagement of oral taste receptor pathways in sensing bacterial-derived signals.

**Figure 1.**
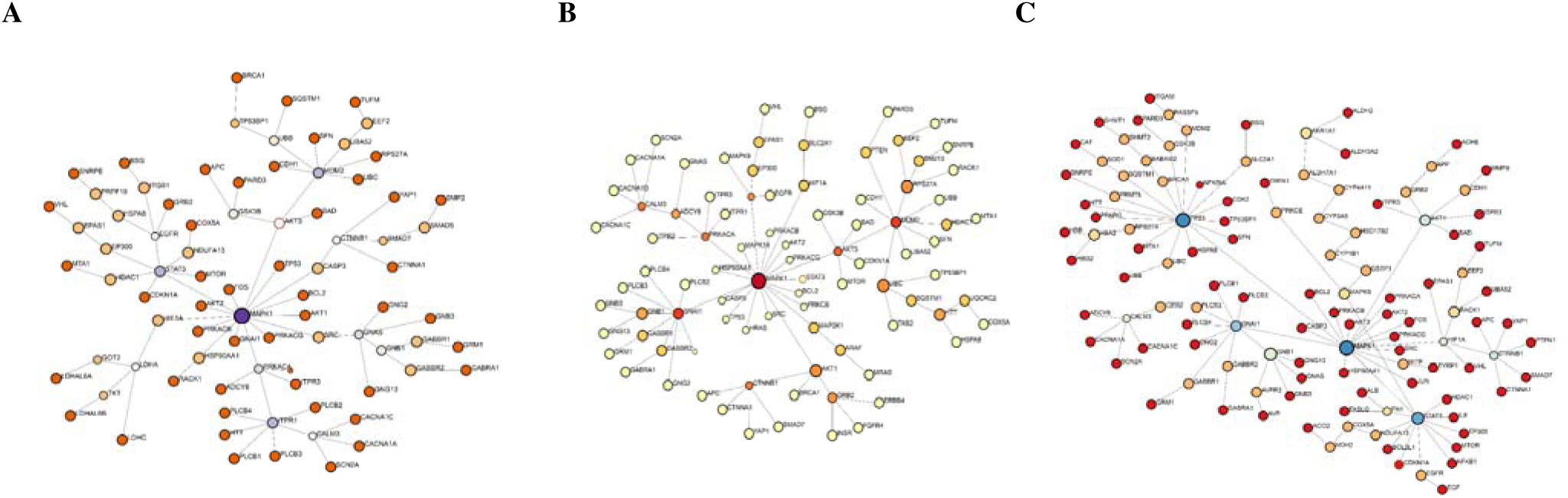
Commensal-associated bacteria preserve taste receptor signaling and sustain cytoprotective autophagy networks in OSCC. KEGG-derived host–bacteria interactomes illustrating signaling architectures associated with (A) *Lactobacillus*, (B) *Lactiplantibacillus plantarum*, and (C) *Streptococcus mitis*. Node size reflects degree centrality and node color indicates betweenness centrality. Across all three taxa, taste receptor–associated GPCR signaling components (GNAS, GNAI1, GNB1/3, GNG2/13, GABBR1/2, GRM1) form a coherent upstream sensory module that couples bacterial detection to intracellular Ca² signaling via canonical intermediates (PLCB1–4, CALM3, ADCY8, ITPR1–3). Downstream integration converges on a MAPK1-centered network that remains functionally distributed rather than forming a rigid inflammatory core. Autophagy regulators (SQSTM1, MTOR, HSPA8) are retained within the core signaling architecture and are positioned alongside checkpoint and tumor-restraining nodes, including TP53, PTEN, CDKN1A, and CTNNB1. Importantly, inflammasome-associated and lytic cell death pathways are not enriched, indicating preservation of autophagic flux and proteostatic balance. Metabolic and stress-response pathways (LDHA, NDUFA13, HIF1A, antioxidant systems) are integrated without dominance, consistent with controlled adaptation rather than oncogenic rewiring. Collectively, these interactomes define a conserved signaling phenotype in which commensal-associated bacteria maintain taste signaling–driven cytoprotective autophagy and checkpoint regulation, supporting a tumor-restraining microenvironment in OSCC.

**Figure 2.**
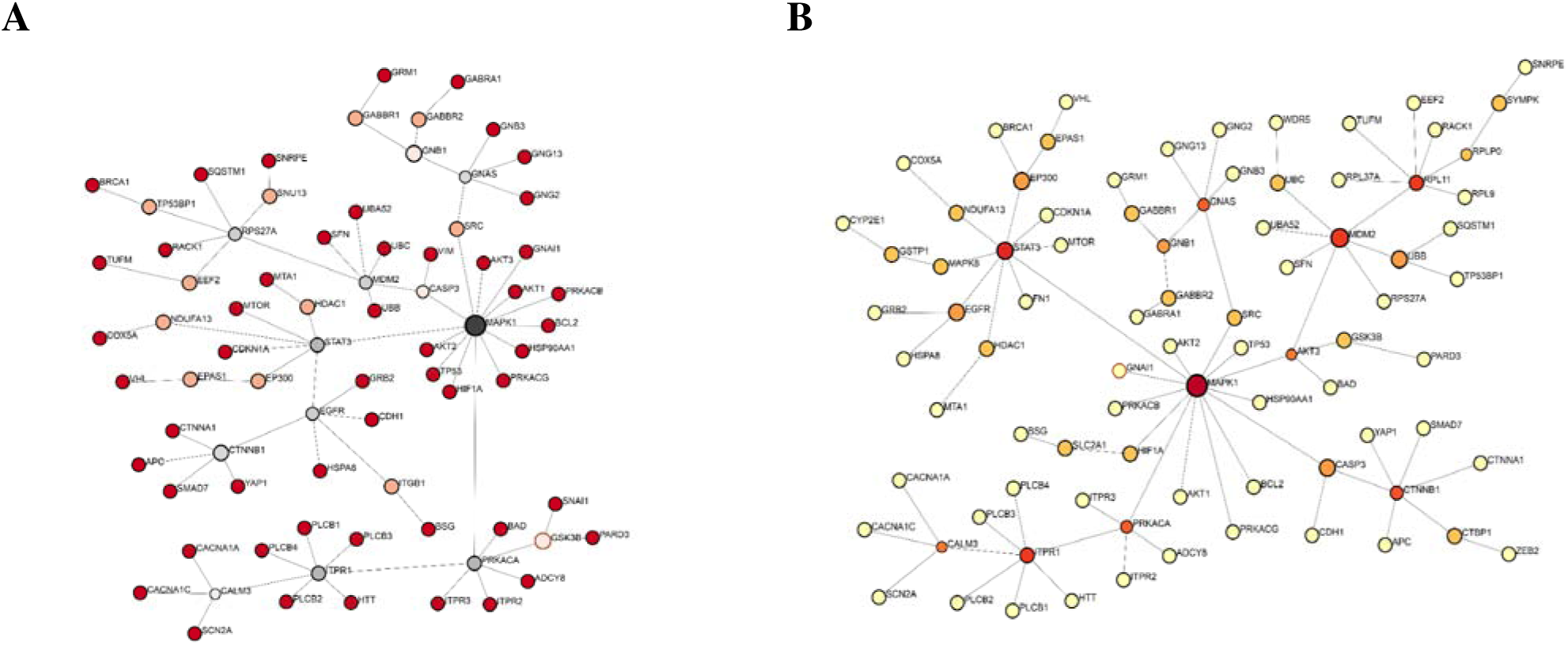
Intermediate bacterial taxa channel taste receptor signaling into stress-adaptive autophagy networks in OSCC. KEGG-derived host–bacteria interactomes illustrating signaling architectures associated with (A) *Capnocytophaga gingivalis* and (B) *Streptococcus gordonii*. Node size reflects degree centrality and node color indicates betweenness centrality. In both taxa, taste receptor–associated GPCR components (GNAS, GNB1/3, GNG2/13, GABBR1/2, GRM1) form coherent upstream sensory modules that connect bacterial detection to intracellular Ca² signaling through canonical intermediates (PLCB1–4, CALM3, ITPR1–3). Downstream signaling converges on MAPK1-centered hubs that integrate chemosensory input with stress and survival pathways, including STAT3, AKT3, PRKACA, and HIF1A. Autophagy regulators (SQSTM1, MTOR, HSPA8) are retained within the core network but occupy an intermediate regulatory position and intersect with checkpoint and remodeling nodes (TP53, MDM2, CTNNB1, HDAC1), rather than aligning with inflammasome or lytic execution pathways. This signaling configuration reflects preserved taste receptor signaling coupled to stress-adaptive autophagy, distinguishing these taxa from both cytoprotective commensals and inflammatory pathogenic profiles. Collectively, these interactomes define a transitional autophagy state associated with context-dependent, weakly pro-tumor– associative behavior in OSCC.

These GPCR inputs consistently propagated through canonical calcium-mobilizing cascades (PLCB1–4, CALM3, ADCY8, ITPR1–3), positioning ITPR1/2 as key relays linking chemosensory activation to intracellular Ca² flux. Downstream, signaling converged on MAPK1-centered networks; however, in contrast to pathogenic profiles, MAPK1 was embedded within a distributed regulatory architecture rather than a rigid inflammatory core. Autophagy-associated components (SQSTM1, MTOR, HSPA8) remained integrated within this network and were aligned with checkpoint and tumor-restraining regulators, including TP53, PTEN, CDKN1A, and CTNNB1, indicating preservation of autophagic flux and proteostatic balance.

Importantly, these networks lacked strong enrichment of inflammasome, NF-κB, or lytic cell death signaling, and instead demonstrated controlled integration of metabolic and stress-response pathways without dominance of proliferative drivers. Collectively, this conserved signaling topology supports a model in which commensal-associated bacteria maintain taste receptor–mediated cytoprotective autophagy and checkpoint regulation, thereby exerting a tumor-restraining effect in OSCC.

### ntermediate bacterial networks define a predicted stress-adaptive autophagy configuration through preserved taste receptor–MAPK1 signaling in OSCC

KEGG-based host–bacteria interactome analysis revealed that intermediate taxa, including *Capnocytophaga gingivalis* and *Streptococcus gordonii*, establish a conserved signaling architecture in which epithelial taste receptor–associated GPCR signaling is preserved but routed into stress-adaptive autophagy rather than fully cytoprotective or inflammatory states (Figure 3). In both taxa, upstream chemosensory modules composed of GNAS, GNB1/3, GNG2/13, and taste-linked receptors (GABBR1/2, GRM1) indicated intact bacterial sensing through oral taste receptor pathways.

**Figure 3.**
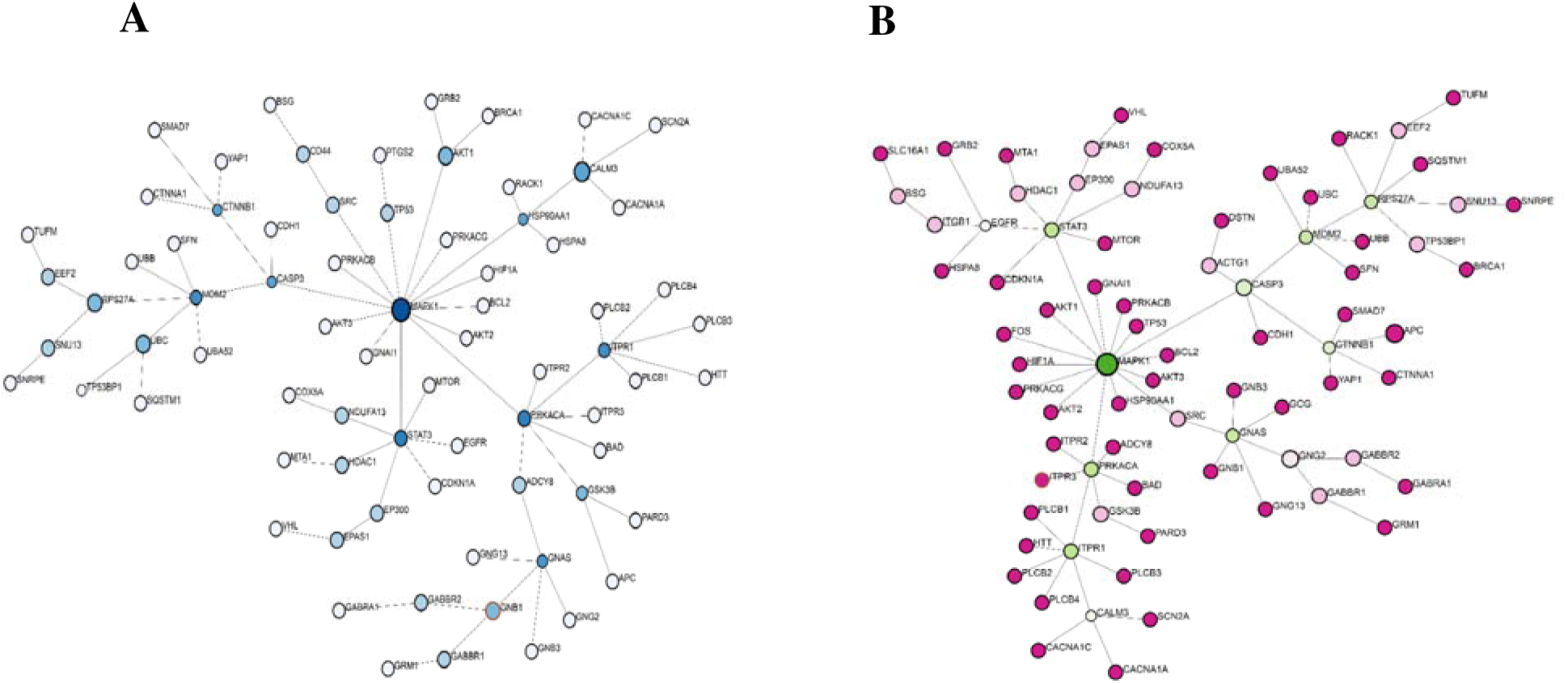
Transition-associated bacteria couple taste receptor signaling to apoptosis-linked autophagy. in OSCC. KEGG-derived host–bacteria interactomes illustrating signaling architectures associated with (A) *Streptococcus anginosus* and (B) *Peptostreptococcus stomatis*. Node size reflects degree centrality and node color indicates betweenness centrality. In both taxa, taste receptor–associated GPCR components (GNAS, GNB1/3, GNG2/13, GABBR1/2, GRM1) form coherent upstream sensory modules that connect bacterial detection to intracellular Ca² signaling through canonical intermediates (PLCB1–4, CALM3, ITPR1–3). Downstream signaling converges on MAPK1-centered hubs that integrate chemosensory input with stress and survival pathways, including STAT3, AKT1/3, PRKACA, and EGFR. Autophagy regulators (SQSTM1, MTOR) are positioned downstream and intersect with apoptotic execution and checkpoint nodes, particularly CASP3, TP53, MDM2, BCL2, and BAD, indicating a shift toward apoptosis-permissive, late-stage autophagy. Unlike strongly pathogenic taxa, these networks do not exhibit dominant inflammasome or lytic signaling but instead reflect a transitional state between adaptive autophagy and inflammatory collapse. Collectively, these interactomes define a MAPK1–CASP3–linked autophagy program associated with context-dependent, pro-tumor–associative signaling in OSCC.

These GPCR inputs propagated through canonical calcium-mobilizing cascades (PLCB1–4, CALM3, ITPR1–3), positioning ITPR1 as a key relay linking chemosensory activation to intracellular Ca² signaling. Downstream integration consistently converged on MAPK1-centered networks; however, unlike commensal-associated profiles, MAPK1 was more tightly coupled to stress and survival signaling modules, including STAT3, AKT3, PRKACA, and HIF1A, indicating a shift toward adaptive stress responses.

Autophagy-associated components (SQSTM1, MTOR, HSPA8) remained embedded within the network but occupied an intermediate regulatory position rather than a purely homeostatic role. Notably, these autophagy nodes intersected with checkpoint and remodeling regulators such as TP53, MDM2, CTNNB1, and HDAC1, without strong enrichment of inflammasome or lytic execution pathways. This distinguishes this group from highly pathogenic taxa while indicating a departure from fully protective autophagy.

Additional engagement of adhesion and epithelial plasticity pathways (ITGB1, CDH1) and moderate integration of metabolic and hypoxia-related signals further support a context-dependent signaling phenotype. Collectively, these findings define a transitional autophagy state in which taste receptor–driven signaling promotes stress-adaptive autophagy and partial oncogenic integration, classifying these taxa as weakly pro-tumor–associative in OSCC.

### Transition-associated bacterial networks predict convergence between autophagy and apoptotic signalling through MAPK1–CASP3 signaling in OSCC

KEGG-based host–bacteria interactome analysis revealed that transition-associated taxa, including *Streptococcus anginosus* and *Peptostreptococcus stomatis*, establish a conserved signaling architecture in which epithelial taste receptor–associated GPCR signaling is preserved but redirected toward apoptosis-coupled autophagy rather than cytoprotective or inflammatory programs (Figure 3). In both taxa, upstream chemosensory modules composed of GNAS, GNB1/3, GNG2/13, and taste-linked receptors (GABBR1/2, GRM1) indicated active engagement of oral taste receptor pathways.

These GPCR inputs propagated through canonical calcium-mobilizing cascades (PLCB1–4, CALM3, ITPR1–3), positioning ITPR1 as a key relay linking bacterial sensing to intracellular Ca² flux. Downstream integration converged on MAPK1-centered networks; however, unlike adaptive profiles, MAPK1 was tightly coupled to apoptotic execution and checkpoint signaling. In both taxa, autophagy-associated components (SQSTM1, MTOR) were retained but positioned downstream and closely aligned with execution nodes, particularly CASP3, as well as checkpoint regulators including TP53, MDM2, BCL2, and BAD.

This network configuration indicates that autophagy is shifted toward an apoptosis-permissive, late-stage response rather than adaptive cytoprotection. Concurrent engagement of oncogenic and remodeling pathways (STAT3, EGFR, CTNNB1, AKT1/3) and adhesion-related components (CD44, ITGB1, CDH1) further supports a transition toward tumor-associated signaling. Notably, these networks lack the strong inflammatory amplification observed in highly pathogenic taxa but demonstrate a clear departure from homeostatic regulation.

Collectively, these findings define an intermediate autophagy state in which taste receptor–driven signaling promotes MAPK1-dependent apoptosis-linked autophagy and stress signaling, classifying these taxa as pro-tumor–associative contributors to OSCC progression.

### Inflammatory and terminal autophagy driven by pathogenic bacteria through MAPK1–STAT3 signaling in OSCC

KEGG-based host–bacteria interactome analysis revealed that pathogenic taxa, including *Prevotella intermedia* and *Fusobacterium nucleatum*, establish a highly centralized signaling architecture in which epithelial taste receptor–associated GPCR signaling is strongly activated but rerouted toward inflammatory and terminal autophagy programs (Figure 4). In both taxa, upstream chemosensory modules composed of GNAI1, GNAS, GNB1/3, GNG2/13, and taste-linked receptors (GABBR1/2, GRM1) indicated robust engagement of oral taste receptor pathways.

**Figure 4.**
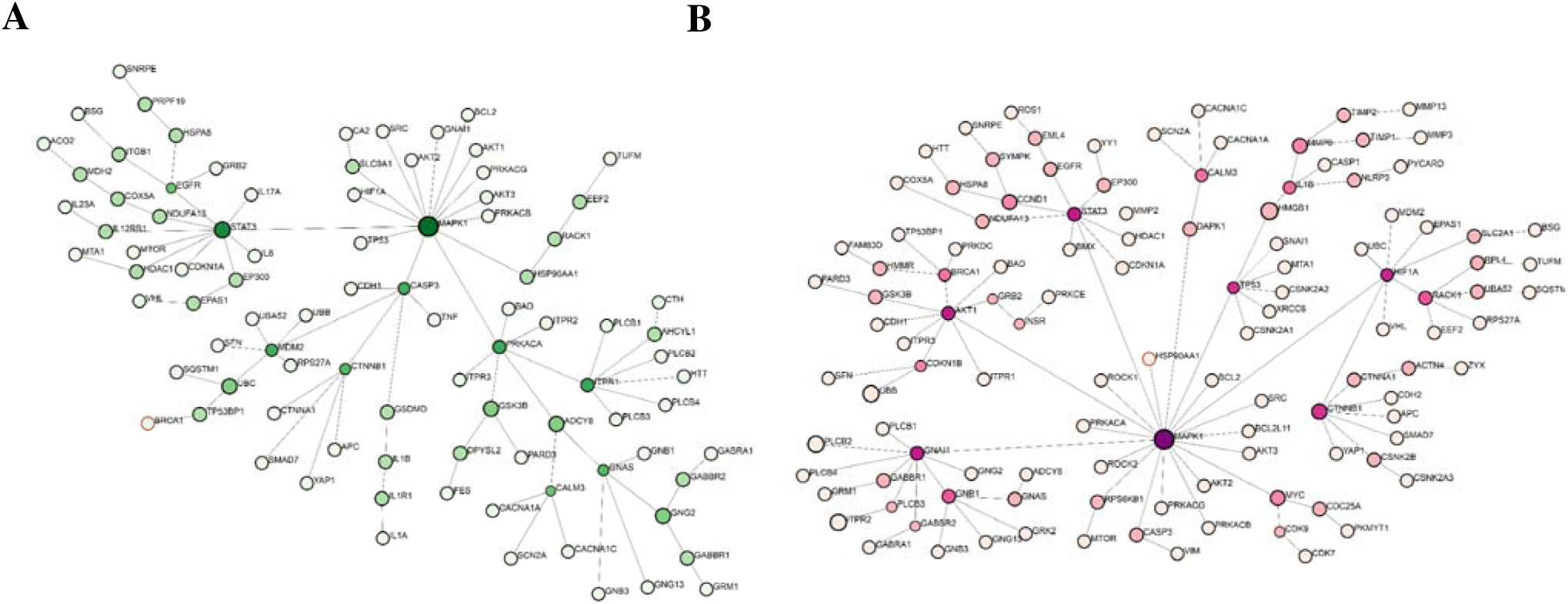
Pathogenic bacteria hijack taste receptor signaling to drive inflammatory and terminal autophagy in OSCC. KEGG-derived host–bacteria interactomes illustrating signaling architectures associated with (A) *Prevotella intermedia* and (B) *Fusobacterium nucleatum*. Node size reflects degree centrality and node color indicates betweenness centrality. In both taxa, taste receptor–associated GPCR components (GNAI1, GNAS, GNB1/3, GNG2/13, GABBR1/2, GRM1) form robust upstream sensory modules that connect bacterial detection to intracellular Ca² signaling through canonical intermediates (PLCB3, CALM3, ADCY8, ITPR1–3). Downstream signaling converges on highly centralized MAPK1-centered hubs that integrate chemosensory input with inflammatory and oncogenic pathways, including STAT3, AKT, PRKACA, EGFR, and HIF1A. Autophagy regulators (SQSTM1, MTOR, DAPK1) are displaced downstream and intersect with inflammasome and lytic execution nodes (CASP3, IL1B, IL1R1, TNF, HMGB1, NLRP3, CASP1), indicating a shift toward inflammation-coupled, terminal autophagy. This signaling configuration reflects hijacking of taste receptor signaling into rigid inflammatory networks, accompanied by activation of proliferative and remodeling pathways (MYC, CCND1, MMP9), and defines a pro-cancer autophagy phenotype associated with OSCC progression.

These GPCR inputs propagated through canonical calcium-mobilizing cascades (PLCB3, CALM3, ADCY8, ITPR1–3), positioning ITPR1 as a key relay linking bacterial sensing to intracellular Ca² flux. Downstream integration converged on densely connected MAPK1-centered networks; however, in contrast to adaptive or transitional profiles, MAPK1 formed a rigid inflammatory core with STAT3, AKT1/3, PRKACA, EGFR, and HIF1A, redirecting taste-derived Ca² signals toward inflammatory amplification and oncogenic signaling.

Autophagy-associated components (SQSTM1, MTOR, DAPK1) were displaced downstream and closely aligned with inflammasome and lytic execution nodes, including CASP3, IL1B, IL1R1, TNF, HMGB1, NLRP3, and CASP1, indicating that autophagy is driven toward an inflammation-coupled, terminal state rather than cytoprotective flux. This shift was accompanied by activation of OSCC-relevant proliferative, metabolic, and remodeling pathways (MYC, CCND1, CDC25A, MMP9, SLC2A1, LDHA) and epithelial plasticity regulators (CTNNB1, YAP1).

Collectively, these findings define a pathogenic autophagy state in which taste receptor–mediated signaling is hijacked to enforce MAPK1–STAT3–driven inflammatory and terminal autophagy programs, classifying these taxa as pro-cancer drivers of OSCC progression.

### The *Porphyromonas gingivalis* network predicts inflammatory dominance and impaired autophagy homeostasis through MAPK1–STAT3–NF-**κ**B signaling in OSCC

KEGG-based host–bacteria interactome analysis revealed that *Porphyromonas gingivalis* establishes a uniquely dysregulated signaling architecture in which epithelial taste receptor–associated GPCR signaling is engaged but rapidly diverted into a highly centralized and dominant inflammatory network, culminating in collapse of autophagy homeostasis (Figure 5). Upstream chemosensory modules composed of GNAI1, GNAS, GNB1/3, and taste-linked receptors (GABBR1/2, GRM1) indicated intact initial bacterial sensing through oral taste receptor pathways.

**Figure 5.**
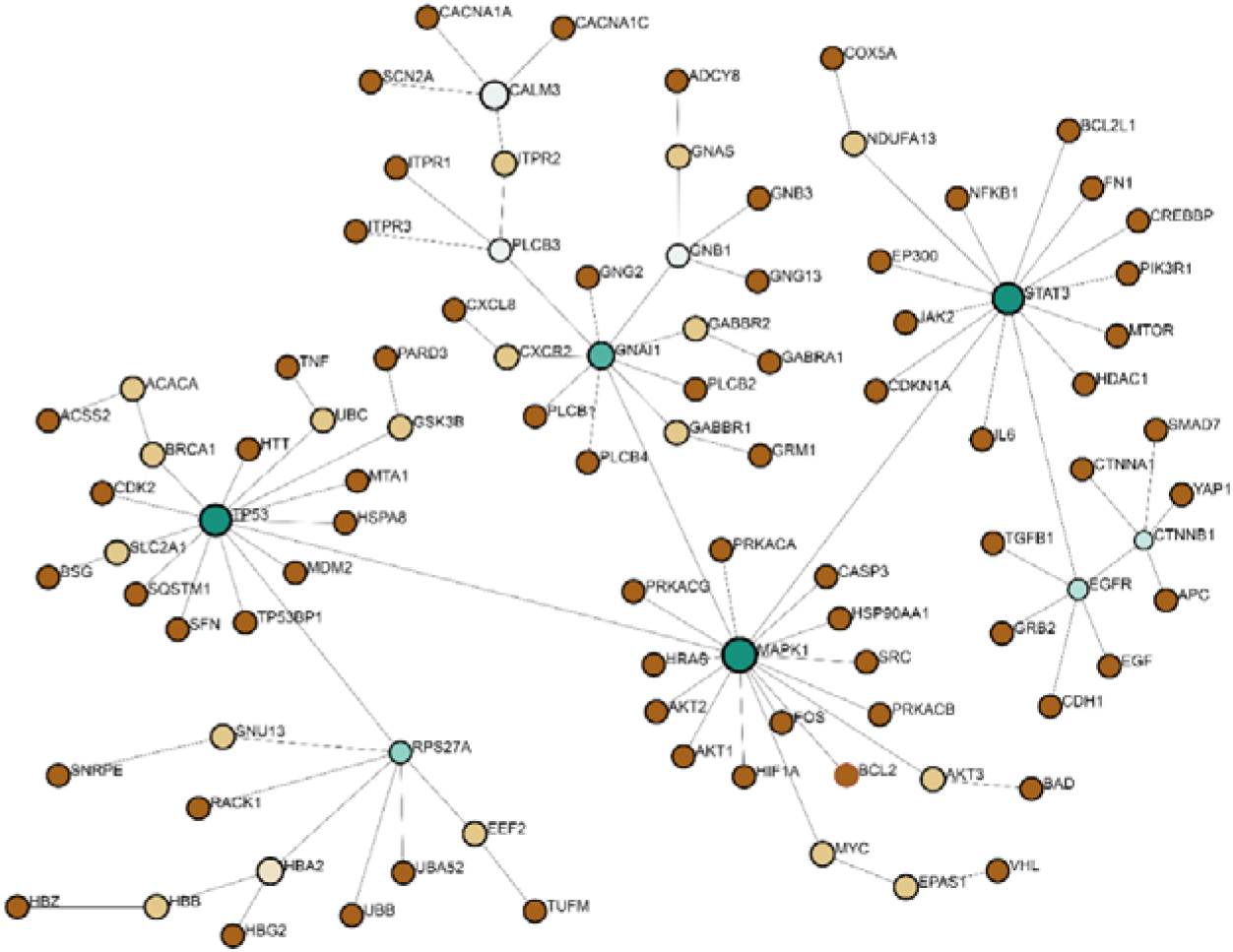
*Porphyromonas gingivalis* hijacks taste receptor signaling to induce autophagy collapse and inflammatory dominance in OSCC. KEGG-derived host–bacteria interactome illustrating signaling pathways associated with *P. gingivalis*. Node size reflects degree centrality and node color indicates betweenness centrality. Taste receptor–associated GPCR components (GNAI1, GNAS, GNB1/3, GABBR1/2, GRM1) connect to Ca²□ signaling intermediates (PLCB3, CALM3, ITPR1–3), establishing upstream bacterial sensing. Downstream signaling converges on a highly centralized MAPK1 hub integrating STAT3, TP53, EGFR, and CTNNB1, forming a rigid inflammatory signaling core. Autophagy regulators (SQSTM1, MTOR) are displaced from central regulatory positions and intersect with inflammatory and execution nodes (CASP3, IL6, CXCL8, NFKB1, TNF), indicating loss of autophagy homeostasis and transition to a dysfunctional, inflammation-coupled state. This network architecture highlights the hijacking of taste receptor signaling into MAPK1–STAT3–NF-κB–dominated pathways, accompanied by activation of metabolic and remodeling programs (MYC, SLC2A1, FN1, TGFB1), defining a terminal autophagy collapse phenotype associated with aggressive OSCC progression.

These GPCR inputs propagated through canonical calcium-mobilizing cascades (PLCB3, CALM3, ITPR1– 3), positioning PLCB3 and ITPR2 as key intermediates linking bacterial detection to intracellular Ca²□ flux. However, in contrast to all preceding groups, this taste receptor–derived signaling was not integrated into adaptive or transitional autophagy networks but instead was rapidly absorbed into a rigid MAPK1-centered signaling core. MAPK1 (Degree 17; Betweenness 3474) formed a dominant hub reinforced by STAT3, TP53, EGFR, and CTNNB1, redirecting signaling toward inflammatory amplification, transcriptional stress, and oncogenic integration. Autophagy-associated components (SQSTM1, MTOR) were displaced to the network periphery and uncoupled from regulatory control, instead aligning with execution and damage-associated signaling nodes, including CASP3, IL6, CXCL8, NFKB1, and TNF. This configuration indicates a loss of functional autophagic flux and transition into a dysregulated, inflammation-coupled state rather than cytoprotective or adaptive autophagy. This collapse of autophagy homeostasis was accompanied by strong activation of OSCC-relevant metabolic and proliferative regulators (MYC, ACACA, SLC2A1, PIK3R1) and epithelial remodeling pathways (FN1, TGFB1, CXCR2), consistent with aggressive tumor-promoting signaling. Collectively, these findings define a terminal pathogenic state in which taste receptor signaling is hijacked to enforce MAPK1–STAT3–NF-κB–driven inflammatory dominance and autophagy collapse, classifying *P. gingivalis* as a key driver of OSCC progression.

Table 3 integrates these findings into a unified framework, demonstrating that bacterial taxa program autophagy along a continuum of signaling states that directly align with OSCC progression.

### Dominant taxon-associated variance and exploratory pathway signature analysis suggest functional patterning of bacterial signaling states in OSCC

Exploratory principal component analysis (PCA) of the full ten-bacteria dataset showed that Fusobacterium nucleatum contributed substantially to the variance observed across the bacterial network signatures (Figure 6A–B). When all ten bacteria were included, the distribution of bacterial network profiles was dominated by two highly separated taxa, Fusobacterium nucleatum and Streptococcus mitis, each occupying distant positions along the principal component axes and disproportionately contributing to overall variance (total variance explained: 51.42%) (Figure 6A). This pattern limited interpretation of the relative relationships among the remaining taxa. To facilitate visualization of the remaining taxa, these two dominant taxa were temporarily excluded and PCA was repeated on the remaining eight bacteria (Figure 6B). This secondary analysis was used for visualization of lower-magnitude variation and was not intended to replace the complete ten-bacteria analysis. This refined analysis suggested apparent separation among the literature-informed pro-cancer, anti-cancer, and intermediate bacterial groups along principal component 1 (PC1; 27.26% variance). Anti-cancer taxa clustered relatively closely, suggesting similar network signatures within the current dataset, whereas pro-cancer taxa displayed greater dispersion, suggesting more heterogeneous network-associated signaling patterns. Intermediate taxa occupied positions between these reference groups within the PCA projection.

**Figure 6.**
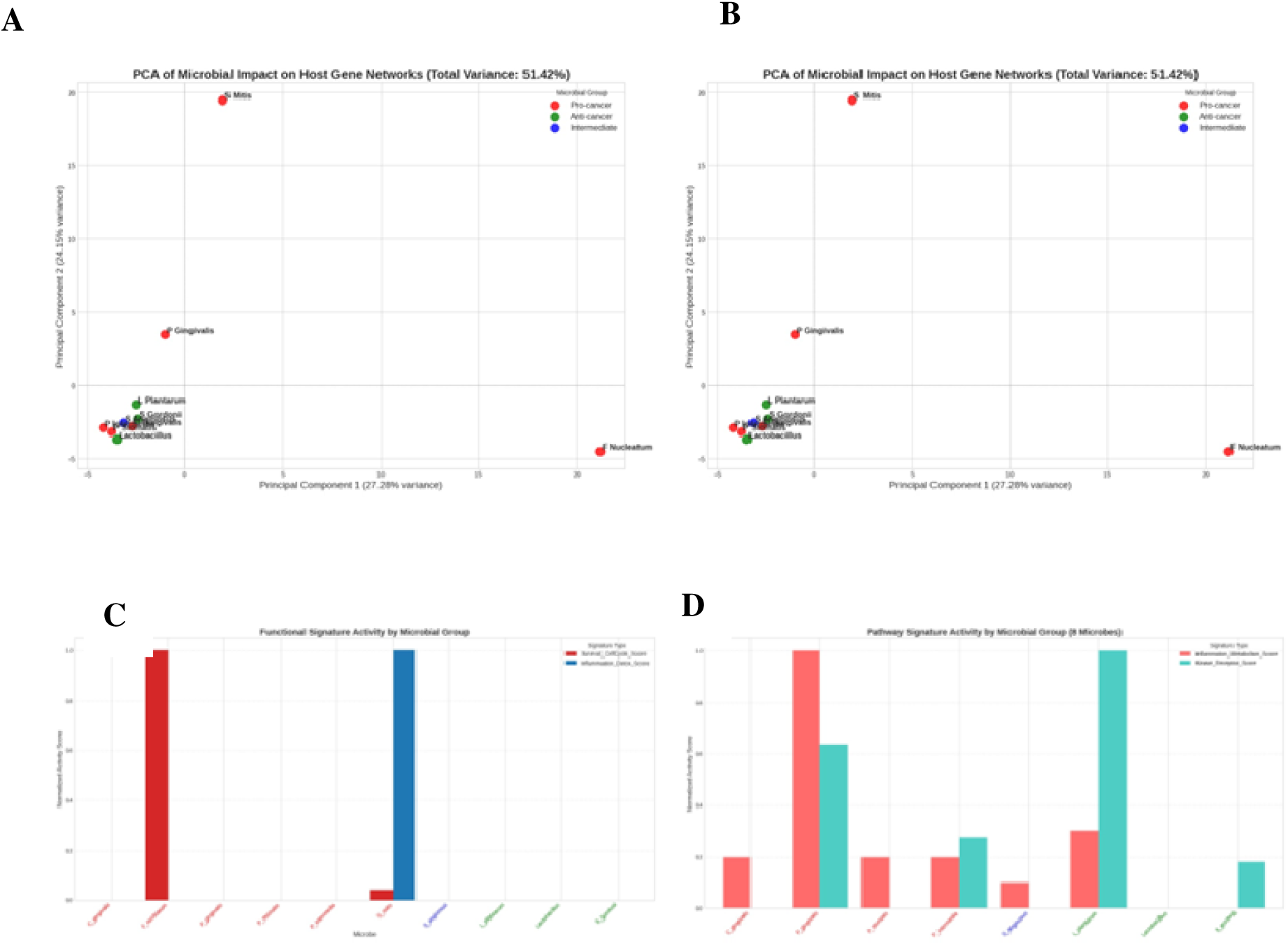
Integrated principal component and pathway signature analysis *suggests functional patterning* of bacterial signaling states in OSCC. (A) Principal component analysis (PCA) including all ten bacterial taxa demonstrates global variance in host gene network signatures, with Fusobacterium nucleatum and Streptococcus mitis acting as *dominant topological profiles that reduce visualization of lower-magnitude relationships among the remaining taxa*. (B) PCA following *temporary exclusion of these dominant profiles for secondary visualization* reveals *apparent separation among the literature-informed* pro-cancer, anti-cancer, and intermediate bacterial groups along principal component 1 (PC1). (C) Group-level pathway signature activity analysis showing *differential representation* of signaling programs, with pro-cancer groups *showing higher inflammatory/tumor-promoting signature scores* and anti-cancer groups *showing greater representation of* homeostatic/checkpoint-associated signatures. (D) Individual taxa-level pathway activity illustrating heterogeneous distribution of *pathway signature scores*, including strongly polarized, mixed, and low-activity bacterial profiles. Together, these analyses suggest that *secondary visualization after accounting for the two dominant profiles reveals lower-magnitude patterns among the remaining taxa*, while pathway activity profiling *supports a potential continuum of network-associated signaling patterns* ranging from cytoprotective to inflammatory and tumor-promoting programs in OSCC. *These findings are exploratory and hypothesis-generating and do not establish independently validated bacterial functional classes or predictive signaling states*.

To further explore the functional basis of this apparent patterning, pathway signature activity analysis was performed at both group and individual taxa levels (Figure 6C–D). At the group level, differences between the two predefined signaling programs were observed (Figure 6C), with pro-cancer bacterial groups exhibiting higher inflammatory/tumor-promoting signature scores, while anti-cancer groups showed greater representation of homeostatic/checkpoint-associated signatures. Intermediate groups displayed comparatively lower scores for both signatures, consistent with their intermediate positioning within this exploratory analysis.

At the individual taxa level, this pattern was further examined (Figure 6D), revealing distinct bacteria with highly skewed pathway signature scores, alongside taxa exhibiting mixed or moderate scores consistent with the literature-informed intermediate reference category. A subset of bacteria displayed minimal representation within both signatures, indicating a lower contribution to the predefined pathway patterns in the current network dataset. Collectively, these analyses suggest that the examined bacterial taxa may exhibit differing pathway-associated network patterns, forming a potential continuum from cytoprotective to inflammatory and tumor-promoting signaling that is broadly consistent with the exploratory network-level patterning observed in OSCC. These findings are hypothesis-generating and do not establish independently validated bacterial functional classes or predictive signaling states.

### Complementary Unsupervised and Statistical Assessment of Bacterial Group Separation

The visual group separation observed in PCA was further assessed using complementary unsupervised and statistical approaches. Hierarchical clustering based on Ward’s linkage method grouped the eight remaining bacterium-specific network profiles into two primary clades that largely corresponded to their literature-informed Pro-cancer and Anti-cancer reference categories (Figure 7A), providing complementary support for the pattern observed in the PCA rather than independent validation of the group structure. To formally assess the stability and non-random structure of the PCA pattern, a permutation test was applied to the PCA results. Comparison of the variance explained by the first two principal components in the real dataset against a null distribution generated by random permutation yielded a p-value of < 0.0001 (Figure 8B), suggesting that the observed separation was unlikely to arise under the specified null distribution. Given the limited number of bacterium-specific networks, this result was interpreted cautiously as supportive evidence of structure within the current dataset rather than proof of generalizability or biological classification. Together, these analyses provide complementary, hypothesis-generating evidence for structured differences among the reconstructed bacterial network profiles.

**Figure 7.**
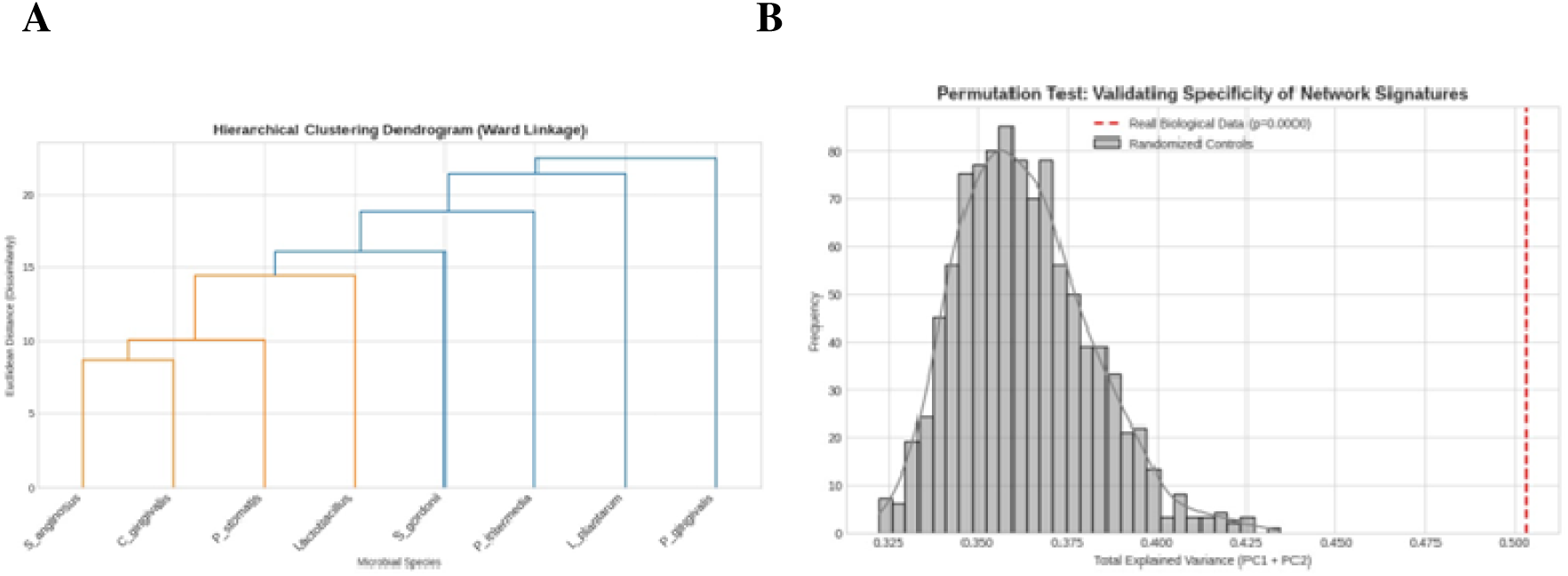
Statistical and unsupervised validation of bacterial grouping. (A) Hierarchical clustering dendrogram showing separation of Pro-cancer and Anti-cancer microbes into distinct clades. (B) Permutation test supporting statistical separation in PCA-derived structure.

**Figure 8.**
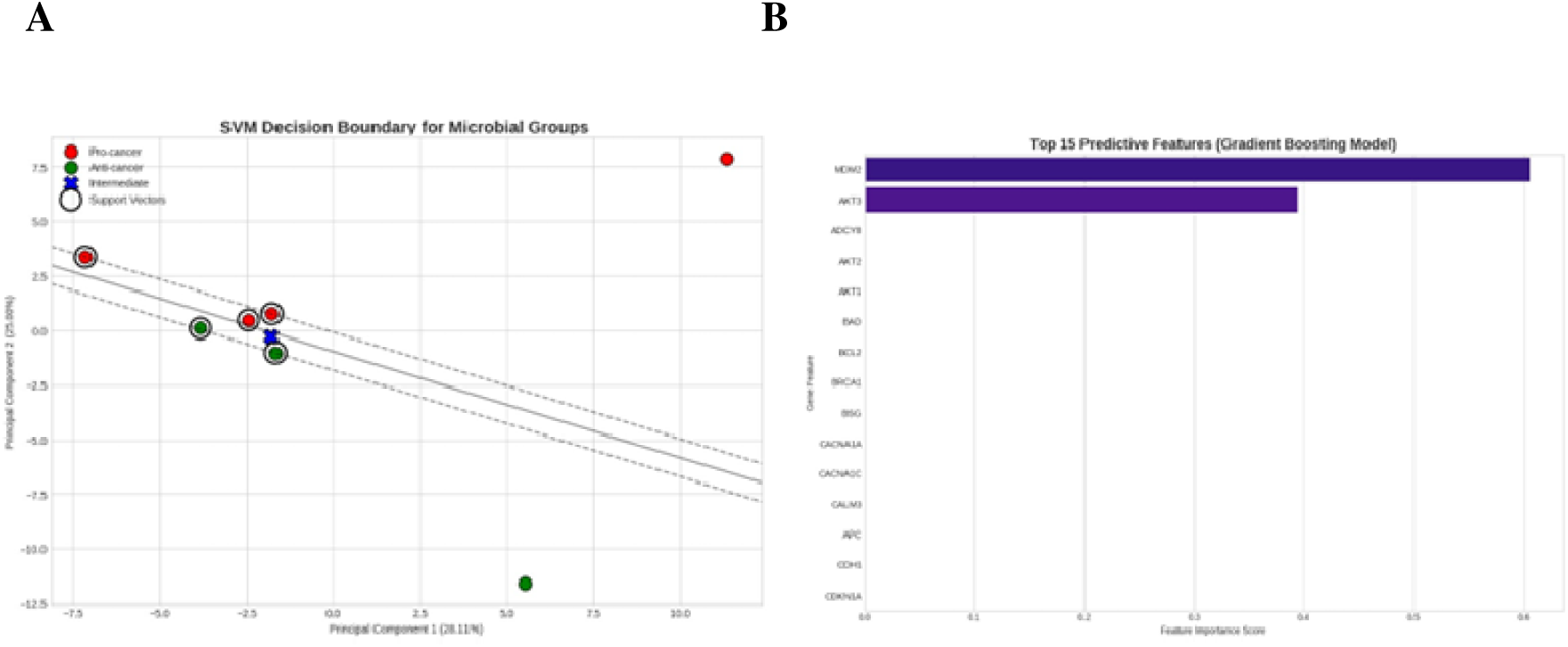
Exploratory supervised analysis of bacterial class separation. (A) Linear SVM decision boundary illustrating separation within the analyzed feature space. (B) Feature importance ranking from the Gradient Boosting Machine identifying MDM2 and AKT3 as dominant model-associated features.

### Exploratory Supervised Analysis Highlights Feature Separability and Candidate Network-Associated Drivers

To further explore whether bacterial *reference categories showed apparent separability* in the network-derived feature space, supervised machine learning models were applied *in an exploratory manner*. A linear support vector machine (SVM) identified a separating hyperplane *between* the Pro-cancer and Anti-cancer bacterial *network profiles* within the analyzed feature space (Figure 8A). This provided a geometric illustration of *apparent category* separability within the exploratory feature space.

The GBM correctly classified the limited held-out observations in this particular split. Because the test set contained only a small number of taxon-level networks, this result was not treated as an estimate of predictive accuracy. Its value was restricted to exploratory prioritization of the network features contributing to separation within the present dataset.However, given the limited sample size, *the small number of test observations, and the literature-informed nature of the reference categories,* these results are interpreted as exploratory rather than as evidence of *stable, generalizable,* predictive performance. Feature importance analysis highlighted the degree centrality of two regulatory genes—MDM2 (importance = 0.61) and AKT3 (importance = 0.39)—as the *highest-contributing features associated with category separation* within the model (Figure 8B). *Because feature-importance estimates can be sensitive to sampling variability in a dataset of this size, MDM2 and AKT3 were interpreted as indicative candidate network-associated contributors rather than definitive biological drivers.* These exploratory results suggest that bacterial modulation of host networks may converge on a limited set of *candidate* regulatory nodes associated with downstream signaling differences.

### Signature Analysis Suggests Differing Functional Patterns

To visualize pathway-level differences associated with group separation, a targeted heatmap was generated using the highest-contributing candidate genes identified by supervised modeling. This analysis revealed differences in the representation of signaling patterns: Pro-cancer microbes showed higher network representation of inflammatory and stress-associated genes, whereas Anti-cancer microbes showed greater representation of checkpoint, kinase-regulatory, and homeostatic signaling modules (Figure 9A). To quantitatively summarize this divergence, a composite Dysbiosis Index was calculated as the difference between inflammatory and kinase-associated pathway activity. This index produced non-overlapping values between the Pro-cancer and Anti-cancer bacterial reference categories within the current dataset (Figure 9B). Despite the limited sample size, the difference approached statistical significance (one-sided Mann–Whitney U test, p = 0.054), providing exploratory support for the potential biological consistency of this network-derived metric. Because the index and its constituent signatures were derived from the same limited network dataset, these findings should be interpreted as hypothesis-generating rather than as validation of a predictive or generalizable dysbiosis measure.

**Figure 9.**
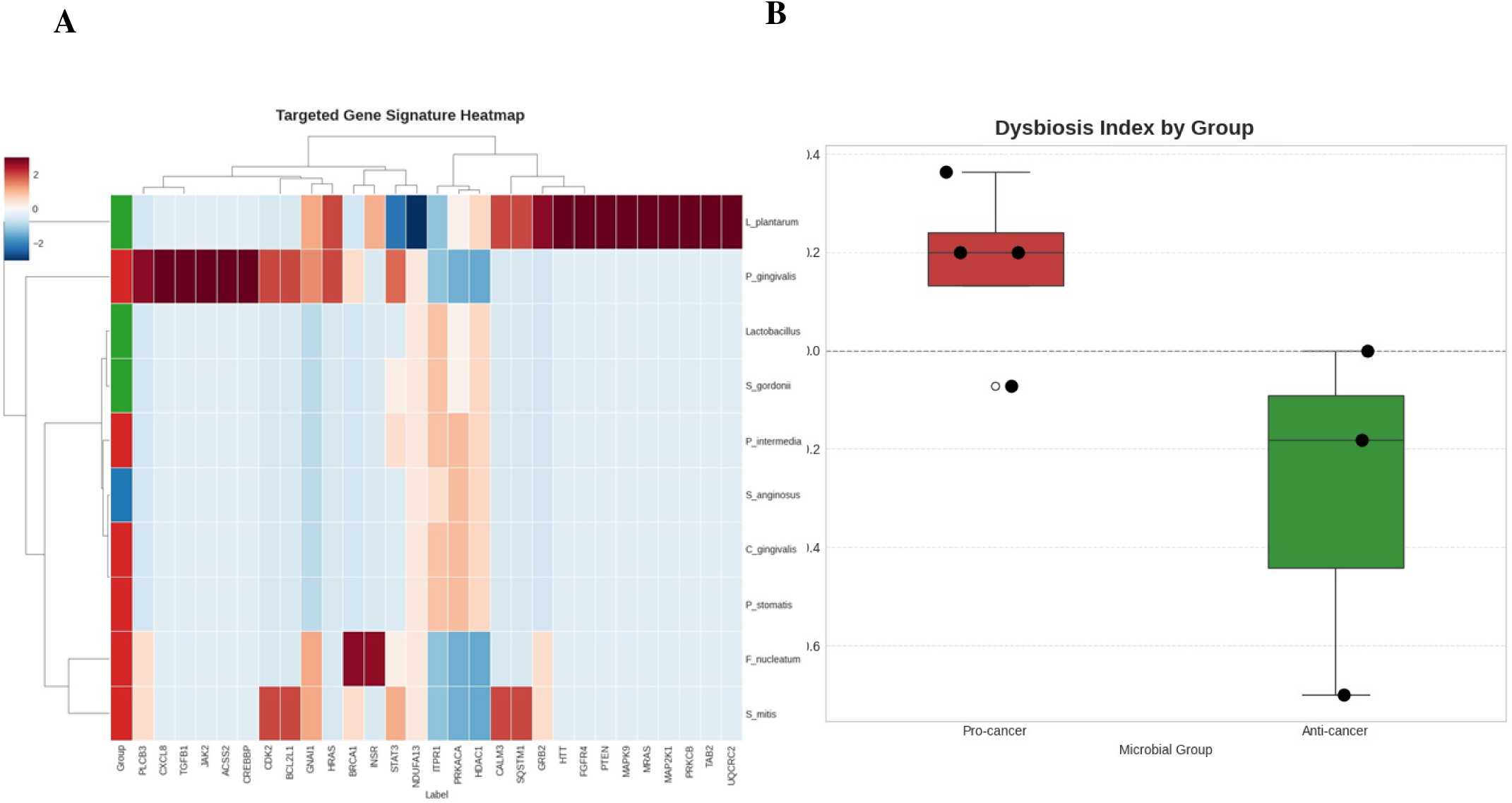
Gene signature heatmap and Dysbiosis Index define functional separation of bacterial signaling states in OSCC. (A) Heatmap of targeted gene signatures illustrating opposing network engagement patterns between pro-cancer (upper cluster) and anti-cancer (lower cluster) bacterial taxa based on key model-associated genes. Distinct clustering highlights differential activation of inflammatory, checkpoint, and stress-response pathways across bacterial groups. (B) Dysbiosis Index analysis quantifying the balance between inflammatory and homeostatic signaling across bacterial taxa. The index demonstrates clear and complete separation between the pro-cancer and anti-cancer groups, supporting functional stratification within the analyzed dataset. Together, these analyses suggest that bacterial taxa segregate into distinct molecular states at both gene and pathway levels, reinforcing the classification of microbial-driven signaling programs in OSCC.

### Docking Studies

To evaluate the binding of the dual-targeting MTDL SG101 to MDM2 and compare it with idasanutlin, docking studies were performed using the crystal structure of human MDM2 in complex with RO5313109 (PDB: 4JRG [61]). Both idasanutlin and SG101 were docked to assess binding poses and interactions within the MDM2 active site.

The MDM2 binding pocket is a deep hydrophobic cleft that accommodates the p53 F19–W23–L26 triad and is primarily defined by residues L53, I57, M58, V89, and I95. As shown in Figure 10, both ligands adopt similar binding orientations and are predominantly stabilized by hydrophobic interactions within this pocket.

**Figure 10.**
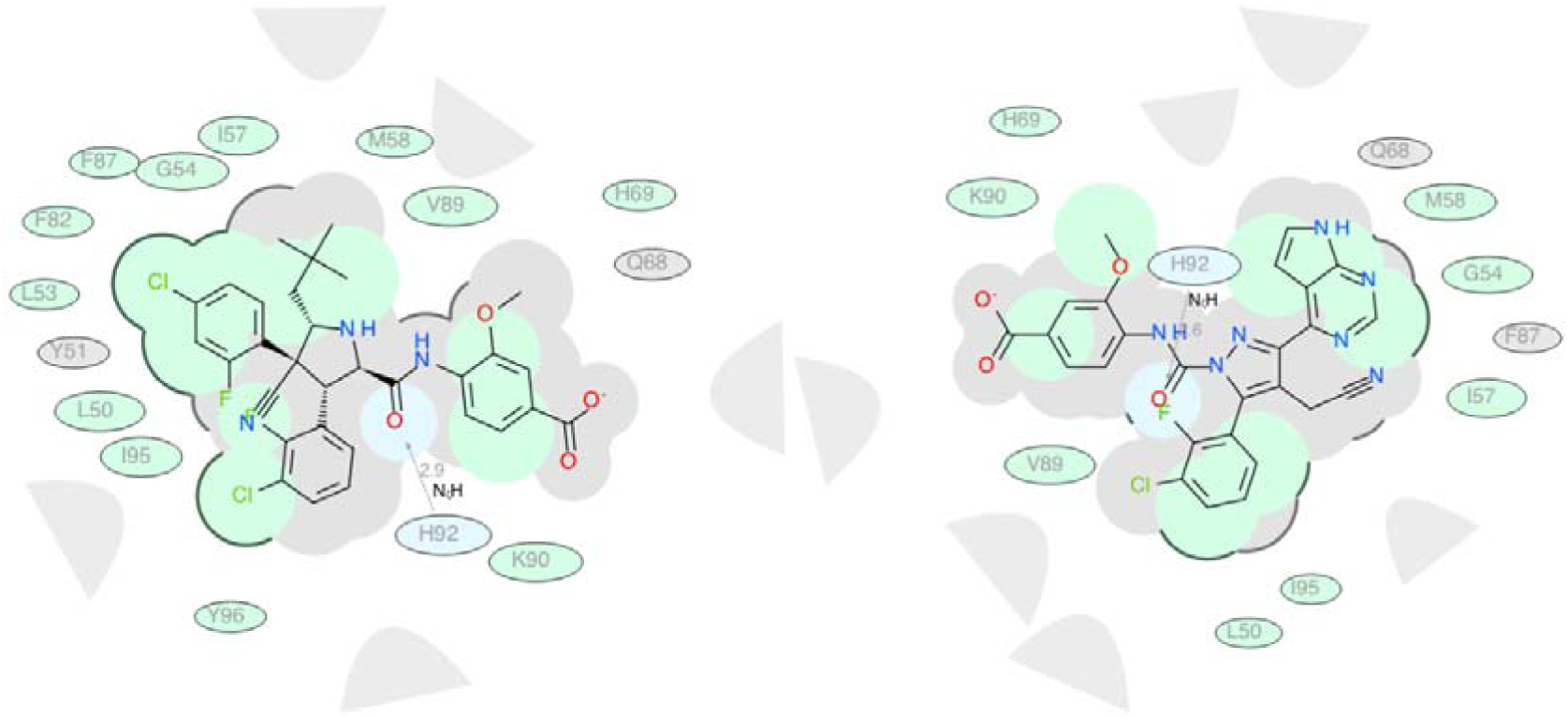
A) The 2D docking pose of idasanutlin and binding interactions within the MDM2 active site (PDB: 4JRG). B) 2D representation of SG101 binding pose within the MDM2 active site (PDB: 4JRG). Ellipticals are representative of amino acid residues. Blue represents hydrogen bonding interactions, green represents hydrophobic interactions, and grey represents van der Wals interactions. Bold outlines around the pocket represent unavailable space, while the absence of outlines represents open space.

For idasanutlin (Figure. 10A), the docking pose reveals a key hydrogen bond between its amide carbonyl and H92 (∼2.9 Å), anchoring the molecule near the polar edge of the binding groove. The halogenated aromatic core is deeply embedded within the hydrophobic cleft, forming extensive contacts with residues L53, I57, M58, V89, I95, F82, and F87. This compact hydrophobic packing, supported by shape complementarity of the halogenated rings, indicates a binding mode dominated by van der Waals interactions. The carboxylate-aromatic terminus remains more solvent-exposed, contributing less to specific binding interactions.

SG101 (Figure 10B) adopts a similar overall orientation, with its central amide linker positioned near the same polar region; however, the hydrogen bond with H92 is longer (2.58 Å), suggesting a weaker interaction. Despite this, SG101 compensates through enhanced hydrophobic interactions, facilitated by its larger fused heteroaromatic system, which increases contact surface area within the pocket. This region interacts with residues M58, G54, I57, F87, and Q68, indicating stabilization through hydrophobic and π-stacking interactions. The halogenated phenyl ring occupies a similar lipophilic region near V89/I95/L50, consistent with conserved binding features between the two ligands. As with idasanutlin, the carboxylate-aromatic moiety of SG101 is oriented toward the pocket entrance and appears largely solvent-exposed, with no strong ionic interaction observed with K90.

Overall, these results suggest that idasanutlin benefits from a stronger H92 hydrogen bond combined with efficient hydrophobic packing, whereas SG101 achieves comparable binding through expanded hydrophobic interactions within the MDM2 pocket. Importantly, Fig. 10B highlights opportunities for further optimization of SG101, including improving linker geometry to strengthen the H92 interaction and modifying the terminal aromatic region to better engage K90. A detailed summary of ligand–residue interactions and distances is provided in Table 4.

**Table 4.** The list of noncovalent interactions of Idasanutlin and SG101 docked in MDM2 (PDB: 4JRG) with the distances from amino acid residues.\.

| Idasanutlin |  |  | SG101 |  |  |
| --- | --- | --- | --- | --- | --- |
| res | type | dist | res | type | dist |
| H92 | hbond | 2.91 | H92 | hbond | 2.58 |
| L50 | hydrophob | 3.04 | L50 | hydrophob | 3.94 |
| L53 | hydrophob | 3.91 | G54 | hydrophob | 3.56 |
| G54 | hydrophob | 3.80 | I57 | hydrophob | 3.61 |
| I57 | hydrophob | 3.35 | M58 | hydrophob | 3.65 |
| M58 | hydrophob | 3.50 | H69 | hydrophob | 4.25 |
| H69 | hydrophob | 4.49 | V89 | hydrophob | 3.64 |
| F82 | hydrophob | 3.48 | K90 | hydrophob | 3.37 |
| F87 | hydrophob | 3.70 | I95 | hydrophob | 3.66 |
| V89 | hydrophob | 3.70 |  |  |  |
| K90 | hydrophob | 3.38 |  |  |  |
| I95 | hydrophob | 2.93 |  |  |  |
| Y96 | hydrophob | 3.67 |  |  |  |

We next evaluated the binding of ruxolitinib and SG101 within the ATP-binding site of the human JAK2 JH1 domain (PDB: 6WTN [62]). The binding pocket consists of a central polar region enabling key hydrogen-bonding interactions, surrounded by hydrophobic subpockets accommodating aromatic and aliphatic groups. As shown in Figure 11, both ligands adopt similar orientations within the polar region, while differing in the extent of polar interactions and hydrophobic occupancy.

**Figure 11.**
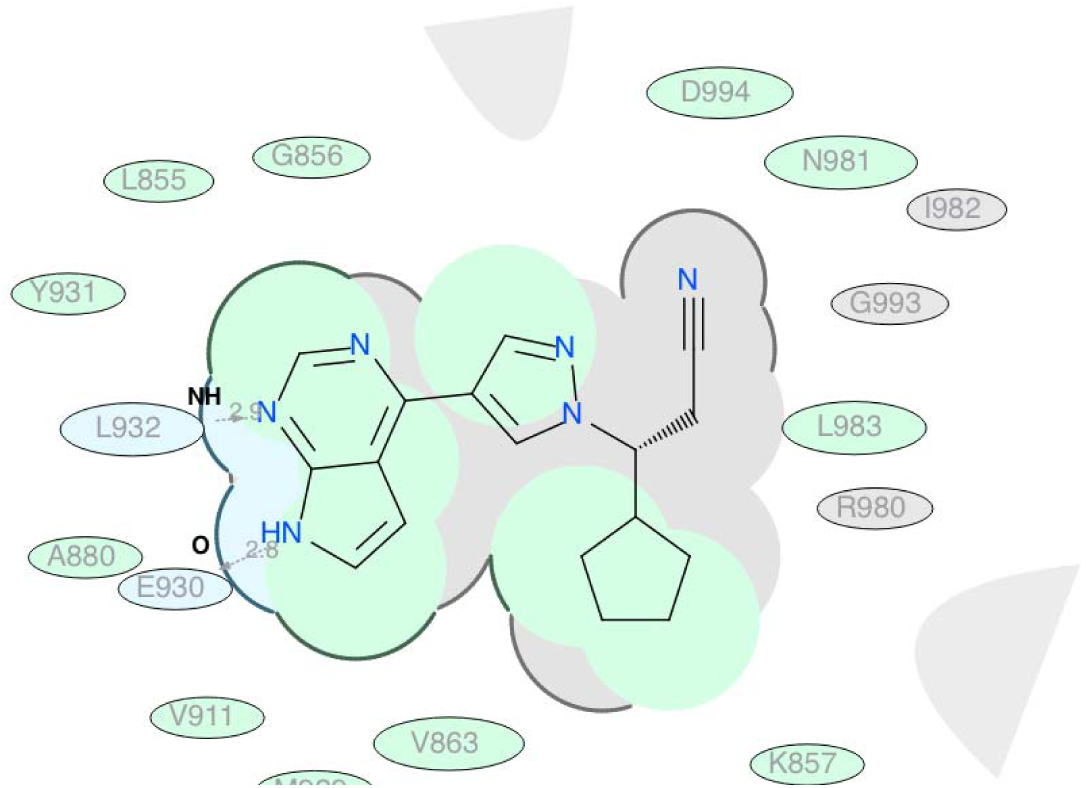
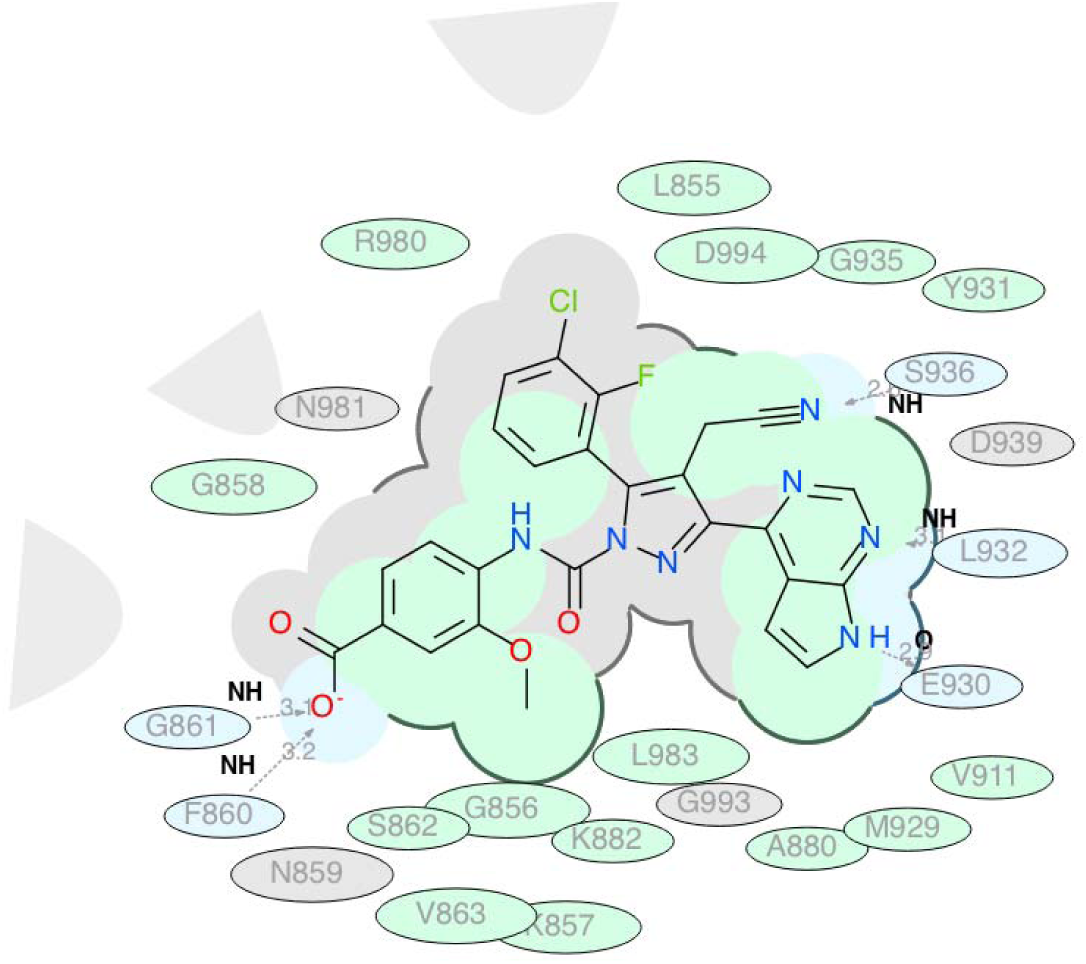
A) The 2D docking pose of ruxolitinib and binding interactions within the ATP kinase active site (PDB: 6WTN). B) 2D representation of SG101 binding pose within the ATP kinase active site (PDB: 6WTN). Ellipticals are representative of amino acid residues. Blue represents hydrogen bonding interactions, green represents hydrophobic interactions, and grey represents van der Wals interactions. Bold outlines around the pocket represent unavailable space, while the absence of outlines represents open space.

Ruxolitinib (Figure 11A) is stabilized by two key hydrogen bonds: one between the pyrimidine nitrogen and L932 (∼2.9 Å), and another between the pyrrole N–H and E930 (∼2.8 Å). These interactions anchor the ligand within the hinge region. Additional stabilization arises from hydrophobic contacts with residues L855, Y931, A880, V911, M929, V863, and G858. The nitrile group extends toward a confined pocket bordered by N981, I982, G993, and D994, suggesting limited tolerance for further substitution in this region.

SG101 (Figure 11B) retains the canonical hinge-binding interactions with L932 (∼3.1 Å) and E930 (∼2.9 Å), while introducing additional polar contacts. Notably, a short hydrogen bond is formed between the nitrile nitrogen and S936 (∼2.2 Å), which is absent in ruxolitinib. Furthermore, the carboxylate-aromatic moiety engages residues G861 and F860 (∼3.1–3.2 Å), providing additional stabilization near the glycine-rich region. SG101 also exhibits expanded hydrophobic interactions with residues L855, Y931, L983, V911, M929, A880, V863, K857, and G856, indicating improved shape complementarity across the ATP-binding site.

Overall, while both ligands share a common binding mode, SG101 demonstrates enhanced interaction density through additional hydrogen bonds and broader hydrophobic engagement. The region extending toward R980 and L855, along with adjacent solvent-exposed space, represents a potential site for further SAR optimization to improve binding affinity without disrupting core interactions. A detailed summary of ligand–residue interactions and distances is provided in Table 5.

**Table 5.** The list of noncovalent interactions of Ruxolitinib and SG101 docked in human JAK2 with the distances from amino acid residues.

| Ruxolitinib |  |  | SG101 |  |  |
| --- | --- | --- | --- | --- | --- |
| res | type | dist | res | type | dist |
| E930 | hbond | 2.80 | F860 | hbond | 3.20 |
| L932 | hbond | 2.94 | G861 | hbond | 3.14 |
| L855 | hydrophob | 3.94 | S936 | hbond | 2.63 |
| G856 | hydrophob | 4.23 | L932 | hbond | 3.13 |
| K857 | hydrophob | 4.08 | E930 | hbond | 2.94 |
| G858 | hydrophob | 3.31 | L855 | hydrophob | 3.72 |
| S862 | hydrophob | 4.05 | G856 | hydrophob | 3.19 |
| V863 | hydrophob | 3.32 | K857 | hydrophob | 3.96 |
| A880 | hydrophob | 3.46 | G858 | hydrophob | 3.40 |
| V911 | hydrophob | 3.75 | S862 | hydrophob | 3.25 |
| M929 | hydrophob | 3.82 | V863 | hydrophob | 3.63 |
| Y931 | hydrophob | 4.12 | A880 | hydrophob | 3.53 |
| N981 | hydrophob | 3.37 | K882 | hydrophob | 4.11 |
| L983 | hydrophob | 3.32 | V911 | hydrophob | 3.79 |
| D994 | hydrophob | 3.88 | M929 | hydrophob | 3.80 |
|  |  |  | Y931 | hydrophob | 4.27 |
|  |  |  | G935 | hydrophob | 4.14 |
|  |  |  | R980 | hydrophob | 3.71 |
|  |  |  | L983 | hydrophob | 3.29 |
|  |  |  | D994 | hydrophob | 3.81 |

## Discussion

*Comprehensive protein-protein interaction network analysis of ten oral bacterial taxa suggests that dysbiosis-associated signals may converge on critical oncogenic signaling hubs linked to OSCC progression. Across all examined networks, bacterial-associated host proteins converged on pathways regulating MAPK signaling, STAT3 activation, and mTOR-mediated autophagy control. This convergence on the MAPK1– STAT3–mTOR axis may represent a candidate mechanism through which pathogenic oral bacteria influence cellular autophagy in a manner potentially favorable to tumor progression. The PI3K/AKT/mTOR pathway is among the most commonly modulated signaling pathways in oral cancer, with mTOR playing an important inhibitory role in regulating autophagy and controlling major cellular and metabolic activities in cancer cells* [7, 63].

Importantly, our results extend beyond a simple binary model of commensal versus pathogenic behavior and instead suggest a potential continuum of bacteria-associated autophagy states, ranging from cytoprotective homeostasis to inflammatory collapse, which may reflect differential routing of taste receptor–mediated signaling across taxa.

A key finding is the contrasting behavior of pathobiont and commensal bacteria across this proposed autophagy continuum. Pathogenic oral taxa can influence inflammatory, prosurvival, and autophagy-related pathways, although their mechanisms and biological effects are species- and context-dependent. *Porphyromonas gingivalis* can modulate NF-κB/IL-6 and PI3K/AKT/mTOR signalling and alter autophagy, producing inflammatory or apoptosis-associated responses in some cellular settings while supporting tumour growth and suppressing apoptosis in OSCC models [7, 64]. *Fusobacterium nucleatum* can promote tumour progression, chemoresistance, and immune suppression through TLR4/MYD88, IL-6/STAT3, and autophagy-related mechanisms [65, 66]. Notably, *F. nucleatum* may activate autophagy to support colorectal cancer chemoresistance or, in a different tumour context, inhibit autophagic flux by disrupting autophagosome–lysosome fusion, demonstrating that its effects on autophagy are not uniform [65, 67]. *Prevotella intermedia* has been linked to tumour invasion and an immunosuppressive tumour microenvironment characterized by increased inflammatory mediators, M2 macrophages, and regulatory T cells, although direct evidence that it regulates tumour-cell autophagy remains limited [68]. *Peptostreptococcus stomatis* can promote colonic tumorigenesis and treatment resistance through ERBB2– MAPK signalling, but its direct involvement in autophagy has not been established [69]. Collectively, cytoprotective, inflammatory, apoptosis-associated, and impaired-flux autophagy should be considered context-dependent responses rather than fixed characteristics of individual bacterial taxa. P. gingivalis increases tumor volume and inhibits OSCC cell apoptosis by promoting cellular autophagy, and is more prevalent in FISH samples from patients with OSCC compared to healthy subjects [7]. These pathobiont-associated networks show autophagy nodes tightly coupled to inflammatory and survival signaling modules, suggesting the potential redirection of homeostatic autophagy toward signaling states associated with tumor progression.

In contrast, commensal organisms such as Lactiplantibacillus plantarum and Lactobacillus spp. demonstrate enhanced connectivity with tumor suppressor-associated signaling, including the GSK3B–APC–CTNNB1 axis, with mTOR centrality notably weak. The beneficial effects of probiotics, such as lactic acid bacteria, *have been associated with activation of* autophagic responses that mediate antitumor effects, underscoring the *potentially* protective role of commensals against malignant progression [7, 70, 71]. These findings are consistent with a *candidate* cytoprotective autophagy phenotype, in which balanced MAPK1 integration with TP53 and PTEN may preserve proteostatic control and *limit* oncogenic escalation.

A particularly important observation is the central role of GPCR and taste receptor-associated signaling in all bacterial networks, with G-protein subunits (GNAS, GNB1/3, GNG2/13) forming *potential* entry points into MAPK and STAT3 signaling pathways. Taste GPCRs possess remarkable structural diversity and normally couple to heterotrimeric G proteins to initiate signal transduction cascades involving activation of adenylyl cyclases, phospholipase C-β2 (PLCB2), production of cAMP, inositol-1,4,5-trisphosphate (IP3), and IP3-dependent Ca² release from the endoplasmic reticulum, with many of these pathways feeding into ERK/STAT3 and mTOR signaling cascades [21]. This *network pattern supports the hypothesis* that bacteria-derived metabolites can engage GPCR/taste receptor pathways, thereby altering calcium and cAMP signaling *and potentially contributing to* oncogenic signaling and dysgeusia in OSCC [72, 73].

Notably, our data suggest that the critical determinant of bacterial impact may not be receptor engagement itself, which is largely preserved across taxa, but rather how downstream Ca² -dependent signaling is routed into distinct MAPK1-centered network configurations associated with different predicted autophagy outcomes.

P. gingivalis exhibits the most densely connected and oncogenic-associated network architecture among all taxa, with STAT3 forming a central hub linking IL6/JAK2 activation and NF-κB-associated components to MAPK1, mTOR, and EGFR. This network density is consistent with experimental evidence indicating that P. gingivalis and F. nucleatum promote cyclin D1 upregulation and apoptosis suppression through altered autophagic flux, thereby prioritizing STAT3-associated signaling for further therapeutic investigation in OSCC. EMT-associated regulators, including CTNNB1, CDH1, and SNAI2, show the strongest network engagement in pathobiont-associated networks, with F. nucleatum having been associated with cellular invasion via crosstalk with E-cadherin/β-catenin signaling and TNFα/NF-κB pathway activation.

Within this framework, our results place these taxa toward one end of the proposed autophagy continuum, where taste receptor signaling may be diverted into MAPK1–STAT3–NF-κB–dominated networks associated with inflammatory amplification and collapse of autophagy homeostasis.

These findings support taste receptor-mediated GPCR signaling as a candidate molecular bridge between dysbiotic bacterial taxa and tumor-promoting autophagy reprogramming, with MAPK1–STAT3–mTOR emerging as a recurrent network-associated regulatory axis. This framework prioritizes potential targets for future investigation in autophagy-aware therapeutic strategies and probiotic-based interventions aimed at restoring homeostatic autophagy in OSCC [70, 74–76].

Building on the network-level identification of convergent signaling hubs, we extended our analysis toward therapeutic translation by integrating structure-based drug design and docking studies. The MAPK1– STAT3–mTOR axis and its upstream regulators, including JAK kinases and MDM2–p53 signaling, represent candidate control points within the bacteria-associated autophagy network identified in this study. These hubs are consistently engaged by pro-cancer reference-category taxa (Table 3), providing a rationale for their exploratory selection as dual-target candidates for pharmacological intervention [77–79].

Docking analysis suggested that the designed MTDL SG101 is capable of engaging both MDM2 and JAK2 binding pockets with favorable predicted interaction profiles. In the MDM2 system (PDB: 4JRG **[50]**), SG101 adopts a predicted binding orientation comparable to idasanutlin, preserving key hydrophobic interactions within the p53-binding cleft while expanding aromatic occupancy to potentially enhance van der Waals stabilization. Although the H92 hydrogen bond is weaker relative to idasanutlin, the increased hydrophobic contact surface suggests possible compensatory binding stabilization. In the JAK2 kinase domain (PDB: 6WTN [62]), SG101 is predicted to retain canonical hinge interactions observed with ruxolitinib while introducing additional hydrogen bonding and broader hydrophobic engagement, suggesting greater predicted interaction density within the ATP-binding pocket [80].

Importantly, ADMET profiling provides preliminary computational support for the translational potential of SG101, demonstrating acceptable predicted drug-like properties, low predicted toxicity, and favorable predicted metabolic stability (Table 2). Together, these findings suggest that SG101 represents a candidate multi-target scaffold with the potential to simultaneously modulate oncogenic signaling and autophagy-related pathways identified in our systems-level analysis.

From a systems perspective, these results suggest that targeting downstream regulatory nodes rather than upstream receptor engagement may provide a potentially more effective strategy to modulate dysbiosis-driven signaling in OSCC.

From a mechanistic perspective, the integration of docking with bacteria–autophagy network analysis provides a hypothesis-generating framework for targeting dysbiosis-driven signaling in OSCC. Rather than focusing on single pathways, this approach may permit investigation of simultaneous intervention at multiple nodes—including GPCR-mediated sensing, calcium signaling, JAK–STAT activation, and autophagy regulation—thereby potentially addressing the redundancy and adaptability of tumor-promoting networks.

This study has several limitations inherent to its hypothesis-generating design. First, the taxon-specific networks were reconstructed from available published bacteria–host interaction evidence and therefore reflect the completeness, context, and potential bias of the existing literature. The ten taxa represent a data-availability-defined cohort and not the full oral microbiome. Second, network connectivity identifies candidate relationships and pathway convergence but does not establish the direction, magnitude, or temporal sequence of biological signalling. Consequently, the proposed autophagy configurations represent network-supported hypotheses rather than direct measurements of autophagic flux. Third, the small number of taxon-level analytical units limits the interpretation of supervised models, classification performance, and feature-importance estimates. These analyses were used to examine internal network structure and prioritize candidate hubs, not to produce a clinically generalizable classifier. Finally, SG101 binding, pharmacokinetic properties, and dual-target activity remain computational predictions. Nevertheless, integration of bacteria-specific networks, taste-associated signalling, autophagy biology, exploratory feature prioritization, and structure-guided ligand design provides a coherent platform for identifying convergent mechanisms and generating testable therapeutic hypotheses in OSCC.Future studies should focus on experimental validation of SG101 binding to MDM2 and JAK kinases, assessment of its impact on autophagy flux, pathway signaling, and tumor cell viability, and evaluation in in vivo OSCC models. Furthermore, expansion of this approach to additional targets within the MAPK1–STAT3–mTOR axis and incorporation of microbiome-modulating strategies may provide a comprehensive framework for developing next-generation, pathway-oriented therapeutics for OSCC.

## CRediT Author Contributions

Hamid Laitifi-Navid and Rui Vitorino: Data curation, bioinformatics analysis, and network modeling. Mahdi Akhavan and Sajjad Aftabi: Machine learning analysis and computational modeling. Stevan Pecic, Manuel Berumen, and Cassandra Yuan: Molecular docking studies and structure-based analysis. Sujatha Peela, SPD Ponamgi, and Robert J. Schroth: Oral health expertise and clinical contextualization of the study. Prashen Chelikani: Conceptual input and critical feedback on taste receptor signaling and OSCC biology. Amir Barzegar Behrooz and Sanaz Vakili: Writing – original draft preparation and data integration. Saeid Ghavami: Conceptualization, supervision, project administration, and integration of autophagy-related mechanisms; writing – review and editing.

